# Transient arousal modulations contribute to resting-state functional connectivity changes associated with head motion parameters

**DOI:** 10.1101/444463

**Authors:** Yameng Gu, Feng Han, Lucas E. Sainburg, Xiao Liu

## Abstract

Correlations of resting-state functional magnetic resonance imaging (rsfMRI) signals are being widely used for assessing functional brain connectivity in health and disease. However, an association was recently observed between rsfMRI connectivity modulations and the head motion parameters and regarded as a causal relationship, which has raised serious concerns about the validity of many rsfMRI findings. Here, we studied the origin of this rsfMRI-motion association and its relationship to arousal modulations. By using a template-matching method to locate arousal-related fMRI changes, we showed that the effects of high motion time points on rsfMRI connectivity are largely due to their significant overlap with arousal-affected time points. The finding suggests that the association between rsfMRI connectivity and the head motion parameters arises from their co-modulations at transient arousal modulations, and this information is critical not only for proper interpretation of motion-associated rsfMRI connectivity changes but also for controlling the potential confounding effects of arousal modulation on rsfMRI metrics.

## Introduction

Resting-state functional magnetic resonance imaging (rsfMRI) signal correlations have been widely used to assess functional brain connectivity and greatly improved our understanding of the intrinsic organization of healthy and diseased brains (Biswal et al. 1995; Fox and Raichle 2007; Zhang and Raichle 2010). However, the validity of a large body of rsfMRI studies was recently challenged by findings about the relationship between head motion and rsfMRI connectivity (Friston et al. 1996; Power et al. 2012, 2014, 2018; Satterthwaite et al. 2012, 2013; van Dijk et al. 2012). Subjects associated with larger head motions during scanning were found to show stronger local rsfMRI connectivity and weaker long-range connectivity (van Dijk et al. 2012). Consistent with this relationship, scrubbing rsfMRI time frames with severe head motion successfully reduced local and recovered long-range brain connectivity (Power et al. 2012, 2018). More recently, the head motion was further suggested to mediate the correlation between the rsfMRI connectivity and behavioral measurements (Siegel et al. 2017). To date, the observed motion-connectivity relationship has been interpreted as a causal relationship with the assumption that the head motion corrupts rsfMRI data and thus functional connectivity derived from it. The relationship is also regarded as a piece of evidence for non-neuronal contributions to the rsfMRI connectivity.

However, the neuronal contribution to this motion-connectivity relationship cannot be completely ruled out. A major motion metric, i.e., differentiated signal variance (DVARS) (Smyser et al. 2010), measures large fMRI signal modulations that are not necessarily caused by the motion, and it often detects brain-wide, synchronized fMRI changes (often called the “global signal”), which have been consistently linked to low vigilance brain states (Fukunaga et al., 2006; He and Liu, 2012; Kiviniemi et al., 2005; Licata et al., 2013; Wong et al., 2016, 2013) and appear to be at least partly neuronal (Schölvinck et al. 2010). This global fMRI signal was recently found to arise from global brain co-activations associated with a neural event of transient arousal modulation showing a characteristic time-frequency signature (Liu et al. 2018). With using multiecho technique to disassociate the neural (R2* contrast) and non-neural (S0 change) fMRI components, a recent study has confirmed that the motion effect on rsfMRI connectivity should be largely attributed to the associated global fMRI signal of R2* contrast that was however assumed to be caused by motion-related respiratory changes (Power et al. 2018). However, the correlation between rsfMRI and respiratory volume (RV) has been found to be highly dependent on the electroencephalogram (EEG) alpha-band power, a major indicator of brain arousal state, and only significant in a sleep-conducive eyes-closed condition but not an eyes-open state, suggesting a significant role of brain arousal in rsfMRI correlations with physiological measures (Yuan et al. 2013). These findings, along with the known association between head motion and sleepiness (Van Den Berg 2006), suggests a possibility that the transient arousal modulations cause both head motion parameters and rsfMRI connectivity changes and thus a spurious relationship between the two. The validity of this hypothesis would reconcile a few puzzling observations about the motion-connectivity relationship. For example, the sensory-dominant pattern of the arousal-related global co-activations (Liu et al. 2018) may explain systematic and divergent changes in local and long-range rsfMRI connectivity associated with head motion. The hemodynamic delay of neurovascular coupling may account for the long-lasting (more than 10 seconds) effects of motion on the rsfMRI connectivity (Power et al. 2014; Byrge and Kennedy 2018). It may also provide a new perspective for understanding the observation that the same amount of head motion causes significant rsfMRI connectivity changes across subjects but not different sessions from the same subject (Zeng et al. 2014).

A clear understanding of the origin of the motion-connectivity relationship is critical for a proper interpretation of findings from a large body of rsfMRI studies. Towards this goal, we use the Human Connectome Project (HCP) data (Van Essen et al. 2013) to examine the role of transient arousal modulations in mediating the association between head motion and rsfMRI connectivity changes. Adapting a template-matching method for fMRI-based arousal quantification (Chang et al. 2016; Falahpour, Chang, et al. 2018), we derived a drowsiness index to locate arousal-related rsfMRI changes. We then demonstrated that the effect of motion-based temporal scrubbing on rsfMRI connectivity can be attributed to arousal-related rsfMRI changes. Lastly, we attempt to understand these correlations between the motion and arousal metrics through their rather complicated temporal relationship based on a trial-based analysis. Overall, the results suggest that the association between head motion parameters and rsfMRI connectivity is not causal but mediated by transient arousal modulations.

## Materials and Methods

### HCP data

The human connectome project (HCP) 500-subject data release was used. The HCP 500-subject data release included 526 subjects who were scanned on a 3T customized Siemens Skyra scanner. We limited our analyses to 469 subjects (age: 29.2 ± 3.5 years, 275 females) who completed all four rsfMRI sessions. Each subject contributed four 15-min sessions on two separate days (two sessions per day). The data were acquired with multiband echo-planar imaging with a temporal resolution of 0.72 seconds and spatial resolution of 2-mm isotropic voxels.

The HCP minimal preprocessing pipelines were applied (Glasser et al. 2013). The fMRI pipelines included the following steps: distortion correction, motion correction, registration to structural data, and conversion to gray-ordinates standard space. Next, the resting state fMRI data were run over the HCP FIX-ICA denoising pipeline to remove artifacts. In addition to the minimal preprocessing pipelines, we smoothed the data temporally (0.001 – 0.1 Hz) and further standardized each voxel’s signal by subtracting the mean and dividing the standard deviation.

### Framewise displacement (FD)

In order to maintain consistency with the previous study on the motion-rsfMRI relationship (Power et al. 2012), we strictly followed its procedures to calculate motion parameters, define regions of interests (ROIs) for rsfMRI connectivity assessment, and estimate the temporal scrubbing effect of different metrics. Framewise displacement (FD) was estimated using a method described previously (Yoo et al. 2005). Specifically, FD was computed as the summing of the absolute values of six differentiated imaging realignment parameters at each time point as follows:

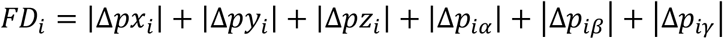

where [*px*_*i*_ *py*_*i*_ *pz*_*i*_] are three translational parameters and [*p*_*iα*_ *p*_*iβ*_ *p*_*iγ*_] are three rotational parameters. |Δ*px*_*i*_| = |*px*_(i−1)_−*px*_*i*_| and the other parameters were calculated in the same way. The three rotational parameters were computed as the displacement of millimeters, which were transformed from degrees on the sphere surface with a rough radius from the cortex to the center of the human head of 50 mm.

### DVARS

DVARS is defined as the rate changes in the fMRI signal averaged over the whole brain at each time point (Smyser et al. 2010). DVARS was computed by applying the root mean square within the whole brain of differentiated fMRI time courses at each time point:

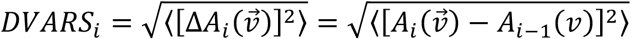

where 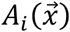 represents the amplitude of BOLD signal at the voxel *v* at the *i*th frame and the angle brackets represent the application of averaging over all of the voxels.

The HCP FIX-ICA denoising pipeline converted DVARS peaks seen in original fMRI signals into sharp DVARS dips, which has been described previously (Glasser et al. 2018), presumably due to an over-correction (**Fig. S1**). In order to maintain consistency with the previous study (Power et al. 2012), we calculated DVARS using fMRI data without applying the FIX-ICA pipeline. We also temporally filtered fMRI data (0.001 – 0.1 Hz) before the DVARS calculations. The DVARS was also calculated with and without the global signal regression (GSR) procedure, but the two versions are very similar to each other (Pearson’s correlation *r* = 0.82 ± 0.17 across 469 subjects, also see an example in **Fig. S1**) and did not result in any major differences in our final results (**Fig. S2)**. We thus reported results using the version with GSR.

### Global signal amplitude (GSA), vigilance index (VI), and drowsiness index (DI)

Global signal was calculated as the mean signal averaged over all gray matter voxels, and the global signal amplitude (GSA) is then defined as the envelop amplitude of the global signal, which was extracted by taking the magnitude of Hilbert-transformed global signals. We adapted a template-matching strategy (Chang et al. 2016) to compute two indices of estimating momentary arousal. Briefly, we took the envelop amplitude of spatial correlations between fMRI co-activation pattern at individual time points and a pre-defined spatial template to estimate the vigilance index (VI) and drowsiness index (DI), and therefore the significant positive and negative correlations with the template are both corresponding to a high value of VI or DI. We used the map of fMRI correlations to EEG vigilance index (Falahpour, Chang, et al. 2018) as the template for computing a vigilance index (VI), and then a global co-activation pattern (Liu et al. 2018) for computing a drowsiness index (DI). Since the envelop amplitude was used to derive these arousal metrics, the large peaks and valleys of the global signal and the spatial correlations to templates would contribute to a high value of these metrics. However, it is worth noting that the large peaks and valleys may represent distinct levels of instantaneous arousal. They often appear as pairs (**Fig. S3**), and may thus represent an arousal drop and a rebound right after it, which together form an event of transient arousal modulation. Therefore, the high DI values are corresponding to periods with frequent transient arousal modulations, which could include transient arousal increase as well.

Both templates are characterized by a sensory-dominant cortical pattern with opposite sign at the dorsomedial thalamus, but the latter is further accompanied by a significant de-activation at the Nucleus Basalis (NB) of the basal forebrain, a subcortical region known to be critical for arousal regulation (Liu et al. 2018). We hoped that this feature may help to improve the sensitivity of detecting transient arousal changes. Overall, these two metrics are strongly correlated with each other (Pearson’s correlation *r* = 0.64 ± 0.18 across 469 subjects), and generate similar results, particularly for the local rsfMRI connectivity (**Fig. S4**), and we thus show the results mostly from the DI. It should be noted that the VI and DI quantify the spatial similarity between individual fMRI time points and pre-defined templates, and are thus not necessarily similar to the GSA. The GSA, VI, and DI were calculated without applying the GSR. Moreover, the GSA, DI and VI were not convolved with the hemodynamic response function. We also calculated a second version of DI, i.e., DI_noSM, with excluding the sensorimotor areas defined by an atlas (Geyer et al. 2000).

### Respiration volume (RV)

Respiration volume (RV) is calculated as the standard deviation of the raw respiratory waveform within a sliding window of 9 TRs (6.48 sec) centered at each time point, with greater robustness to noise than respiration volume per unit time (RVT) (Chang et al. 2009).

### Regions of interest

All fMRI time courses used in this study were extracted from a set of 264 regions of interest (ROIs), which are spheres of 10 mm diameter centered on the coordinates given by the previous study (Power et al. 2011). The rsfMRI correlations among the 264 ROIs were calculated after the global signal regression procedure to be consistent with the previous study (Power et al. 2012). We also compared the temporal scrubbing effect of DVARS and DI using the correlations of 264 ROIs calculated without global signal regression procedure (**Fig. S5**). The connectivity changes in **Fig. S5** were dominated by the global effect, and the distinct local and long-range changes were not visible.

### Relating arousal-related behavioral measures to the motion and arousal metrics

Among a total of 295 behavioral, demographic, and physiological measures released with the HCP data set, we selected and correlated 20 items from the Pittsburgh Sleep Questionnaire (PSQ) with mean FD, DVARS, GSA and DI averaged over four sessions of each subject. Bed times after midnight were treated as the original bed times plus 24 hours. Note that the subject who slept around 3pm was regarded as an outlier and removed during analysis.

### Temporal scrubbing procedure

The first and last 20 time points from FD, DVARS, GSA, DI and RV time courses were removed from the subsequent analyses since they showed some edge effects from the temporal filtering. For each of the metrics, a temporal mask was generated to mark the 25% time points of all the subjects showing the highest values, and this threshold was chosen based on the previous study (Power et al. 2012). The threshold of scrubbing marking top 25% values for FD, DVARS, GSA, and DI is 0.203 mm, 0.0084, 0.1179, and 0.123 respectively. The analysis was also performed with using two other thresholds, i.e., 10% and 40%. A control mask was also created by circularly shifting the FD mask by 600 time points. We then calculated rsfMRI Pearson’s correlations of the 264 pre-defined ROIs for unscrubbed (*r*) and scrubbed (*r*′) data for these masks. Their difference (Δ*r* = *r*′ − *r*) was calculated and then averaged across subjects. The differences (Δ*r*) were plotted as a function of Euclidean distance between the ROIs. The rsfMRI connectivity changes for ROI pairs with a Euclidean distance of 13 ~ 49 mm and of 125 ~ 161 mm were averaged to represent local and long-range connectivity changes respectively. The two ranges of distance were chosen arbitrarily to cover the regimes showing most prominent rsfMRI connectivity changes.

In addition, to estimate the metric-specific scrubbing effect with excluding the arousal-related changes, we also calculated a metric-specific mask for any metrics with excluding its overlaps with the DI mask (see **Fig. S6** with two examples using this original approach). Given that the metric-specific masks actually excluded the top 25% DI points, their DI values are significantly lower than that of all the time points and similar to that of the bottom 75% DI points, and they thus would also act as a mask of low DI values. To better control the DI effect, we estimated the metric-specific scrubbing effect in a different way. Specifically, for each metric-specific mask, we semi-randomly found the same amount of time points with similar DI values as these metric-specific time points and similar metric values as all the time points (**Fig. S7**). This group of control time points were found as follows. For each time point covered by the metric-specific mask, we randomly picked up a time point from all those that have a similar DI value with absolute difference less than 0.001. Then, the metric-specific scrubbing effect was defined as the difference of rsfMRI connectivity with removing the metric-specific time points and this control group of time points. In this way, we can truly separate the metric-specific scrubbing effect with a good control of the DI effect. We estimated the DI-specific scrubbing effect with respect to a metric in a similar way (**Fig. S7**). A control group of time points were found to match the metric value of the DI-specific mask and the DI value of all the time points, and the DI-specific scrubbing effect was then defined as the difference in rsfMRI connectivity calculated with removing the DI-specific time points and this group of control time points.

We also tested the effect of various pre-processing steps on the final results. Temporal scrubbing effects of DI and DVARS were examined using the DVARS calculated without applying global signal regression (**Fig. S2**). Temporal scrubbing effects of DI and DVARS were examined with scrubbing different amounts of time points. The 25% top values in DVARS or DI were scrubbed in **Fig. 4**. We also performed the same analysis using two other thresholds, i.e., 10% (**Fig. S8**) and 40% (**Fig. S9**). Temporal scrubbing effects of DI and DVARS were also examined with skipping certain pre-processing steps before computing rsfMRI connectivity. The temporal filtering step was skipped for the functional connectivity calculation in **Fig. S10**. The FIX-ICA procedure was skipped for the functional connectivity calculation in **Fig. S11**. Both the temporal filtering step and FIX-ICA procedure were skipped for the functional connectivity calculation in **Fig. S12**. The global signal regression procedure was skipped for the functional connectivity calculation in **Fig. S5**. Temporal scrubbing effects of DI and DVARS were also examined by convolving DVARS with a canonical hemodynamic response function from SPM using its default settings (the delay of response: 6 seconds; the delay of undershoot: 16 seconds; the ratio of response to undershoot: 6; the length of kernel: 32 seconds) in **Fig. S13**. Temporal scrubbing effects of RV and DI were examined in **Fig. 5**. We also convolved RV with a respiratory response function (Birn et al. 2008) and examined its scrubbing effect with DI in **Fig. S14**. Temporal scrubbing effects of FD and DI were examined in **Fig. S15**. We also convolved FD with a hemodynamic response function and examined its scrubbing effect with DI in **Fig. S16**.

### Temporal dynamics of various metrics around FD spikes

For this part of the analysis, the mean of FD, DVARS, DI, and RV time courses were removed for each subject to reduce inter-subject variation in the baselines. FD spikes were located by finding peaks 3 standard deviations above the mean of each FD time course. Segments of various metrics centering on the detected FD spikes (± 28.8 seconds) were extracted and then sorted according to the DI values at 9.36 second (13 TR), where a proportion of DI segments show consistent peaks. After sorting, the top and bottom 10% segments (~400 each) were averaged respectively and their difference was also calculated.

## Results

We analyzed HCP rsfMRI data of 469 subjects who completed all four resting-state sessions in two different days. Four indices were derived from the rsfMRI signals. Framewise displacement (FD) (Yoo et al. 2005) and DVARS (Smyser et al. 2010) have both been used for assessing head motion, but they may present different information since the former is derived directly from image alignment parameters whereas the latter quantifies the amplitude of fMRI changes between consecutive time points. The global rsfMRI signal has been consistently shown to inversely related to brain vigilance state (Fukunaga et al., 2006; He and Liu, 2012; Kiviniemi et al., 2005; Licata et al., 2013; Wong et al., 2016, 2013), we thus took its envelope amplitude, referred to as the global signal amplitude (GSA), as an approximate estimate for instantaneous arousal level. Large peaks of the global rsfMRI signal have been shown to be associated with a neurophysiological event indicative of transient arousal modulation, and also show a sensory-dominant co-activation pattern with specific de-activations at the dorsomedial thalamus (DMT) and the Nucleus Basalis (NB) at the basal forebrain, the subcortical regions known to get involved in arousal regulation (**Fig. 1A**) (Liu et al. 2018). To incorporate this spatial information, we adapted the idea of a template-matching approach (Chang et al. 2016), correlated this global co-activation template with individual rsfMRI volumes, and then took its envelop amplitude as an index to estimate arousal fluctuations (**Fig. 1B**). Because the global fMRI co-activations appear most frequently at intermediate states of vigilance (Liu et al. 2018), such as drowsy state and light sleep, we named the fourth metric as the drowsiness index (DI) (See Methods for details about how to derive these four metrics).

**Figure 1.**
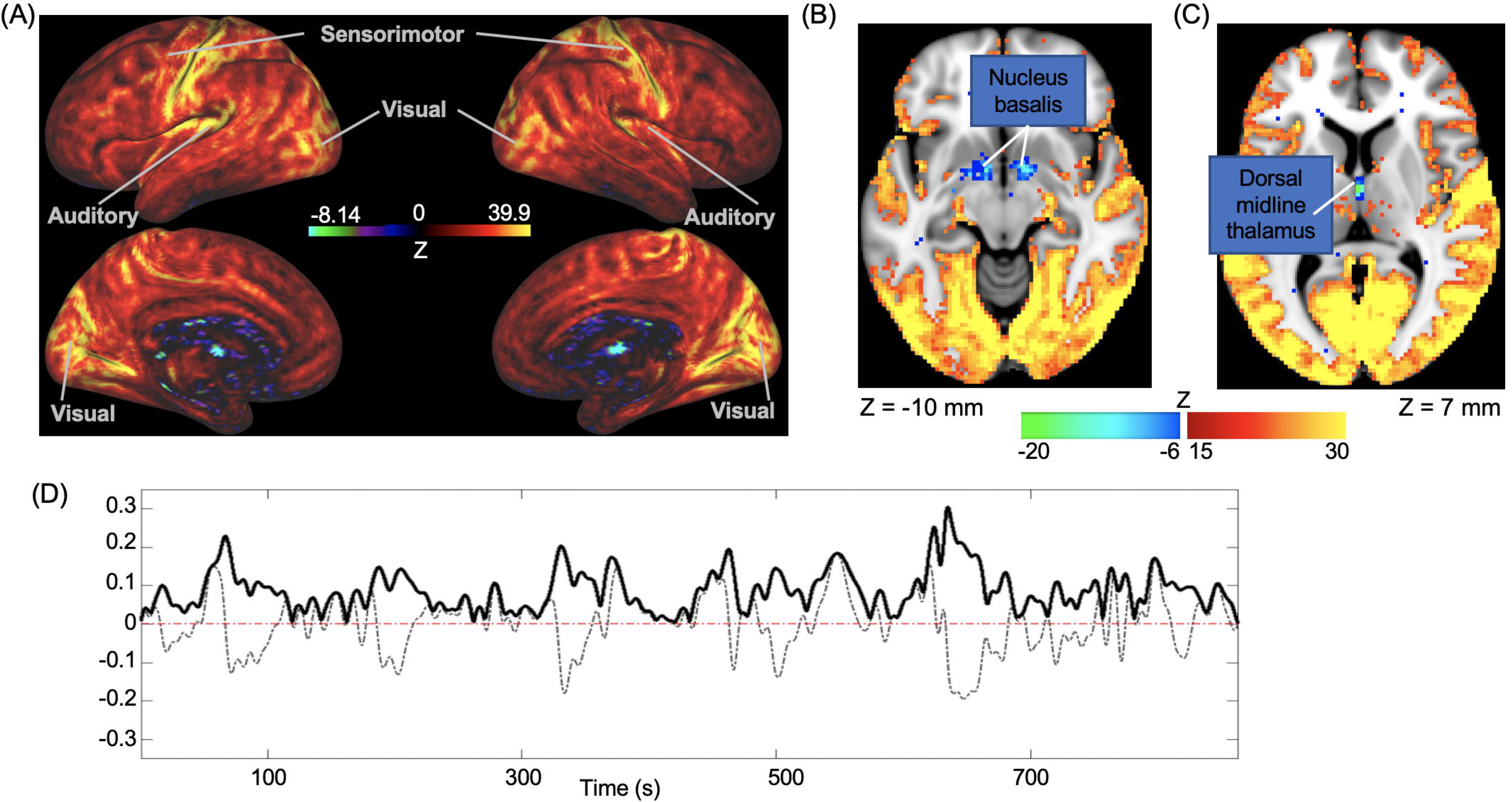
Drowsiness index (DI) derived from a representative subject. (A) The global fMRI co-activation pattern, which has been shown to be associated with an event of transient arousal modulation, is used as a template for deriving DI. Sensorimotor, visual, and auditory regions show a larger signal increase than other cortical regions and the nucleus basalis (Z = −10) (B) and the dorsal midline thalamus (Z = 7) (C) show signal decrease. (D) An example of the DI time course of a representative subject. DI is defined as the envelope amplitude (solid black) of the spatial correlations (dashed gray) between the global co-activation pattern (A) and individual fMRI volumes.

### Motion and arousal metrics are related to arousal

All four metrics showed a tendency of increasing amplitude over the course of rsfMRI scanning. This effect was much stronger for the arousal indices, i.e., the GSA and DI, but also present for the motion metrics, i.e., the FD and DVARS (**Fig. 2A**). This temporal trend likely indicates an increasing probability for subjects becoming drowsy or falling asleep, which is consistent with a previous finding that a significant proportion of subjects fell asleep within 3 minutes into the resting-state scanning (Tagliazucchi and Laufs 2014).

**Figure 2.**
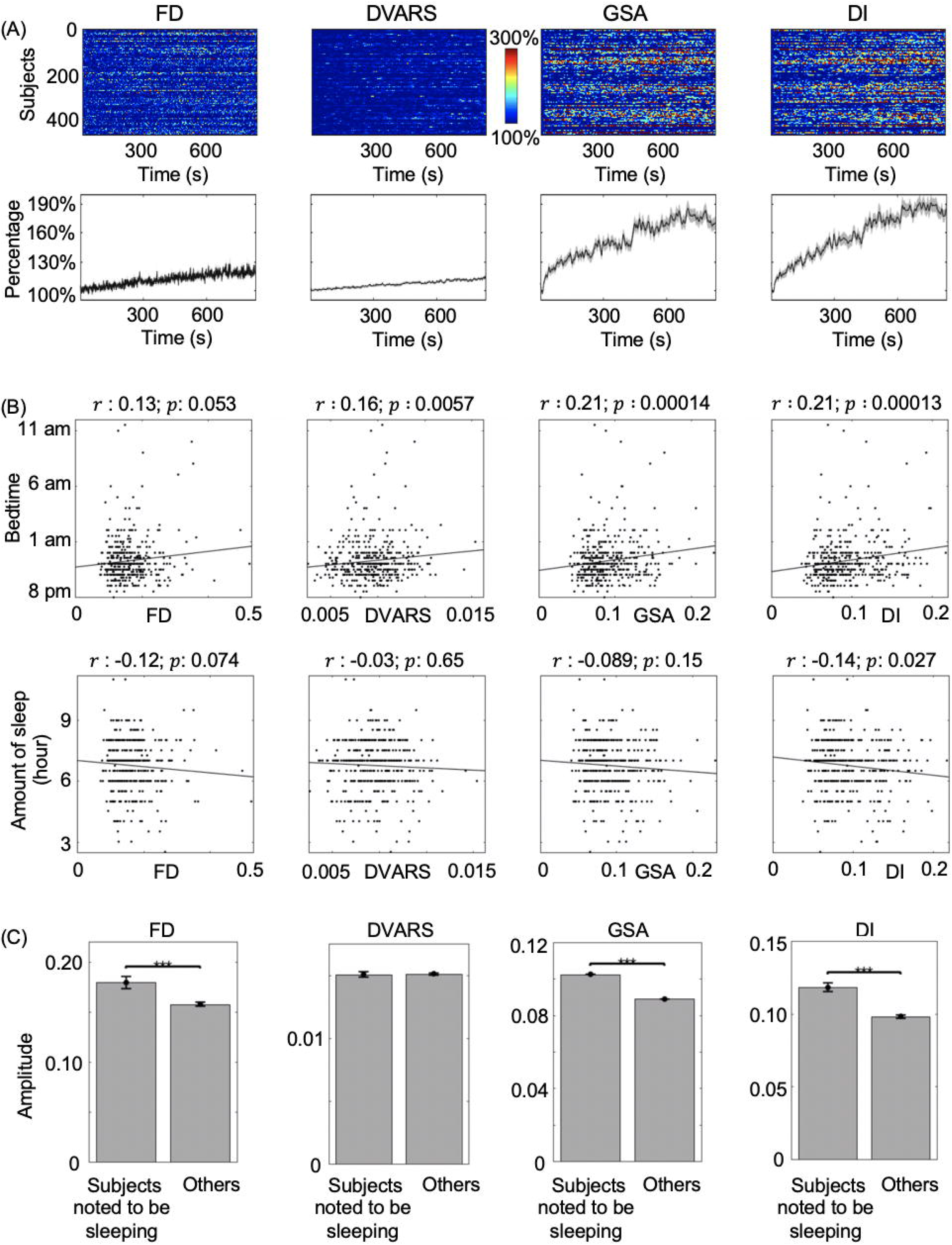
Motion and arousal metrics are related to brain arousal. (A) The amplitude of FD, DVARS, GSA, and DI increases over the course of resting-state scanning. Time courses of the four metrics, which were normalized with respect to their initial values during the first 14.4 seconds (20 TRs), are shown for each of 469 subjects (top panels), as well as their averages with the shaded region representing area within 1 standard error of the mean (SEM) (bottom panels). (B) Cross-subject Pearson’s correlations between the mean FD, DVARS, GSA, and DI scores and two PSQI items, i.e., Bedtime and Amount of Sleep. Significant correlations were found between these two arousal-related measures and the DI scores, with Pearson’s correlation coefficients and corresponding False Discovery Rate (FDR) corrected values being shown on the top. (C) 72 subjects, who were noted to be sleeping during the resting-state scanning sessions, show significantly higher DI score compared with the other subjects who were not. The error bar represents the SEM. The asterisks represent the level of significance: ***, *p* ≤ 0.001, two sample *t*-test, df = 467.

In addition to the intra-subject temporal trend, we also examined the relationships between these metrics and arousal-related behavioral measures across subjects. We correlated the individual’s DI score, defined as the mean DI averaged over all 4 sessions, with the 20 Pittsburgh Sleep Questionnaire (PSQ) items provided by the HCP (**Table S1**). FD and DI are significantly correlated with multiple PSQ items. The items correlated with DI appear to be related to the sleep propensity of subjects (**Fig. 2B** and **Table 1**). The subjects who go to bed later (Pearson’s correlation *r =* 0.19, *p* = 9.4×10^−4^, degree of freedom (df) = 467, same as below) and have less amount of sleep (*r = −*0.14, *p* = 0.027, df = 467) showed significantly higher DI scores. In contrast, FD is significantly correlated with the items reflecting the comfort level of sleeping (**Table S1**). The subjects who reported to snore (*r =* 0.23, *p* = 3×10^−5^, df = 467), cannot breathe comfortably (*r =* 0.16, *p* = 0.007, df = 467) or felt too hot (*r =* 0.14, *p* = 0.027, df = 467) during sleep are associated with significantly higher FD values. In addition, the DI score is significantly higher (*p* = 1.5×10^−10^, Hedges’ *g* = 0.66, two sample t-test *t* = 6.70, df = 467, **Fig. 2C**) in a group of subjects who were noted to be sleeping by the scanning operators during the resting-state fMRI scanning (Glasser et al. 2018). Overall, both the intra- and inter-subject analyses indicate that there is a close relationship between the motion/arousal metrics, particularly DI, and the arousal-related measures.

**Table 1.** Pearson’s correlation of three representative alert-related behavioral measures with FD DVARS, GSA and DI (df: 467; FDR-corrected *p* values less than 0.05 are highlighted)

| Behavioral measures | FD |  | DVARS |  | GSA |  | DI |  |
| --- | --- | --- | --- | --- | --- | --- | --- | --- |
|  | Pearson | FDR- | Pearson | FDR- | Pearson | FDR- | Pearson | FDR- |
| | $r$ | corrected $p$ values | $r$ | corrected $p$ values | $r$ | corrected $p$ values | $r$ | corrected $p$ values |
| PSQI_Score | 0.14 | **0.027** | 0.060 | 0.30 | 0.045 | 0.47 | 0.063 | 0.30 |
| Bed time | 0.13 | 0.053 | 0.16 | **0.0057** | 0.21 | **0.00014** | 0.21 | **0.00013** |
| Amount of sleep | -0.12 | 0.074 | -0.030 | 0.65 | -0.089 | 0.15 | -0.14 | **0.027** |

**Position of Table 1:** around the second paragraph of the section *Motion and arousal metrics are related to arousal* in the result part.

### Temporal scrubbing effects on rsfMRI connectivity based on different metrics

Scrubbing time points with a high value of motion metrics has become a commonly used approach to alleviate the influence of the motion on rsfMRI connectivity (Power et al. 2012, 2014, 2018). Here we compared the effects of temporal scrubbing on rsfMRI connectivity based on the four metrics. We assessed the whole-brain rsfMRI connectivity using Pearson’s correlation coefficients between fMRI time courses of each possible pair of 264 regions of interest (ROIs) (Power et al. 2011), then estimated temporal scrubbing effects by the connectivity difference before and after scrubbing time points with the top 25% values in a specific metric.

Consistent with the previous findings (Power et al. 2012, 2014, 2018), the temporal scrubbing based on FD (one-sample *t* = 3.59, *p* = 5.43×10^−4^, Cohen’s *d* = 0.23, df = 468 for local connectivity; *t* = 0.99, *p* = 0.34, Cohen’s *d* = 0.064, df = 468 for long-range connectivity; same as below; **Fig. 3A**) and DVARS (*t* = 11.26, *p* = 0, *d* = 0.73, df = 468 for local connectivity; *t* = 0.82, *p* = 0.36, *d* = 0.054, df = 468 for long-range connectivity; **Fig. 3B**) decreased local rsfMRI connectivity as compared with a control case to remove the same amount of irrelevant time points (**Fig. 3E**). Nevertheless, the scrubbing based on FD, which were computed directly from image alignment parameters (Yoo et al. 2005), showed a much smaller (*t* = 10; *p* = 0; *d* = 0.81; df = 468; FD vs DVARS based scrubbing effect on the local connectivity) effect than the DVARS-based scrubbing. This could be due to the fact that DVARS (Smyser et al. 2010) is, by definition, sensitive to large fMRI changes that are not necessarily caused by the head motion. Scrubbing time points of high GSA values produced a significant effect for the local (*t* = 15.37; *p* = 0; *d* = 1.02; df = 468) but not for long-range connectivity (*t* = 1.85; *p* = 0.08; *d* = 0.12; df = 468), which are at a similar level as the DVARS-based scrubbing effect. The largest changes in both local and long-range rsfMRI connectivity were observed with excluding the top 25% DI time points (**Fig. 3D**), which is significantly larger than both DVARS-based (*t* = 5.52, *p* = 0, *d* = 0.35, df = 468 for local connectivity; *t* = 5.69, *p* = 0, *d* = 0.39, df = 468 for long-range connectivity) and GSA-based (*t* = 4.32, *p* = 3.5×10^−5^, *d* = 0.22, df = 468 for local connectivity; *t* = 4.01, *p* = 1.1×10^−4^, *d* = 0.28, df = 468 for long-range connectivity) scrubbing (**Fig. 3F**). In summary, the arousal-related fMRI changes, flagged by high DI values, affect rsfMRI connectivity in a similar way as the head motion parameters does but to a much larger extent.

**Figure 3.**
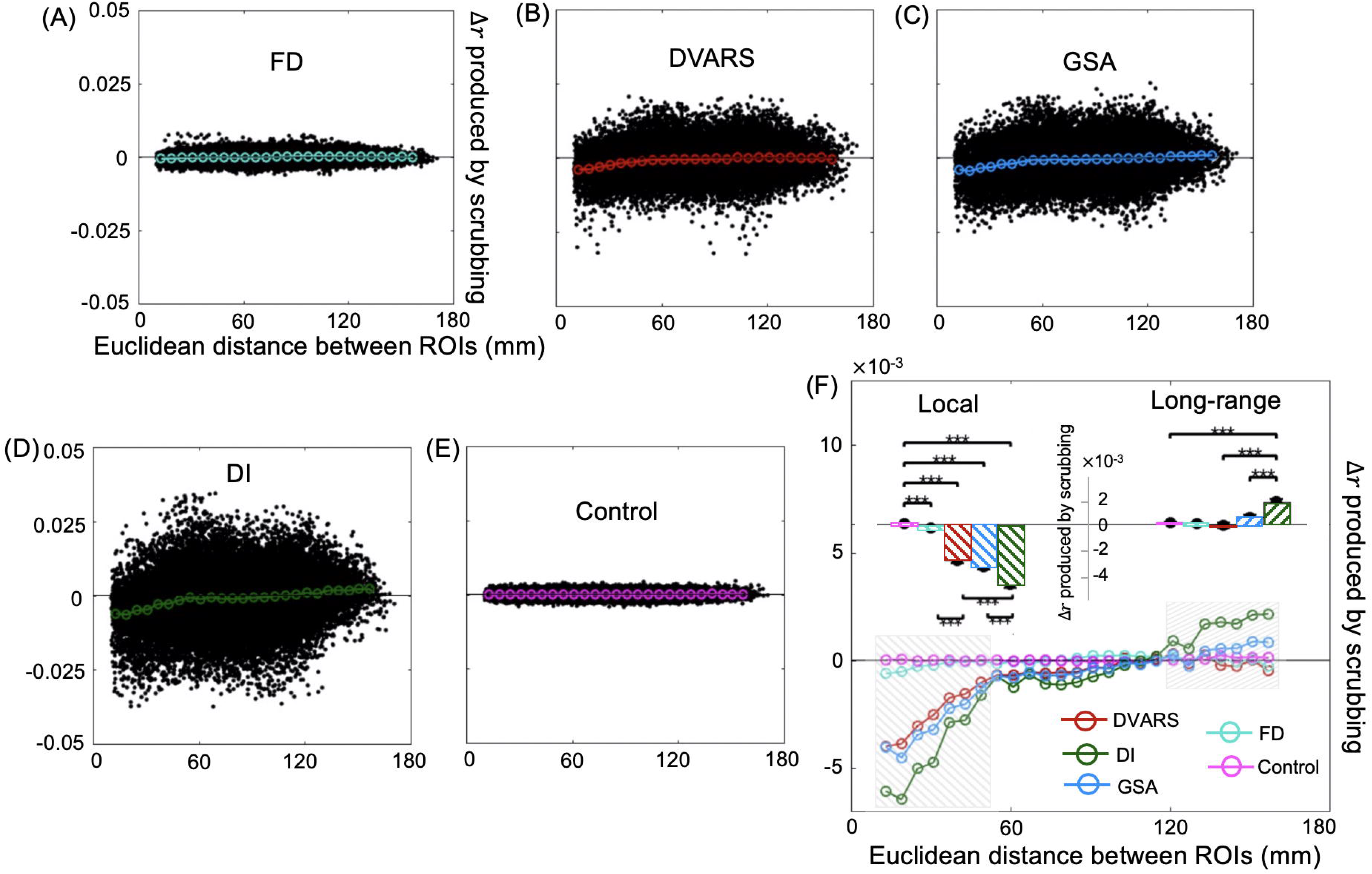
Effects of temporal scrubbing on resting-state fMRI connectivity. Removing time points with the top 25% values in FD (A), DAVRS (B), GSA (C), and DI (D) reduced the local but increased the long-range rsfMRI correlations between the 264 pre-defined brain ROIs, whereas temporal scrubbing using a control mask, which is created by circularly shifting the FD mask by 600 time points, produced no effects on rsfMRI connectivity (E). A comparison of temporal scrubbing based on different metrics (F) suggests that the DI-based scrubbing has the largest effect on rsfMRI connectivity. Black dots in (A-E) represent the rsfMRI connectivity difference before and after scrubbing with respect to their distance for each pair of ROIs across all subjects. The colored circles in (A)-(E) represent the averaged connectivity changes every 6 mm and are summarized into (F). The local (ROI pairs with a distance between 13 and 49 mm, gray box) and long-range (ROI pairs with a distance between 125 and 161 mm, gray box) rsfMRI connectivity changes are summarized as a bar plot and shown as an inset in the top region in (F). The error bar represents SEM across subjects. The asterisks represent the level of significance: ***, *p* ≤ 0.001 with df = 468.

We then examined metric-specific scrubbing effects since significant overlaps are expected between scrubbed time points based on different metrics. Indeed, 39% of scrubbed (i.e., 9.81% of the total) time points are the same for the DVARS- and DI-based scrubbing, and this value is significantly higher (*p* = 0; permutation test, *n* = 10,000) than the one expected (6.25% of the total) with assuming these two metrics are completely independent. After excluding these overlapped time points from scrubbing, i.e., retaining them for rsfMRI connectivity assessments, the DVARS-specific scrubbing produced marginal significant changes on either local (*t* = 3.19; *p* = 0.00746; *d* = 0.21; df = 468) or long-range (*t* = 3.73; *p* = 0.001339; *d* = 0.24; df = 468) rsfMRI connectivity, but the effect of DI-specific scrubbing remained large and significant for local (*t* = 12.80; *p* = 0; *d* = 0.83; df = 468; **Fig. 4**) and long-range connectivity (*t* = 5.86; *p* = 0; *d* = 0.38; df = 468; **Fig. 4**). A similar result was obtained for the pair of FD and DI (**Fig. S15**). We repeated the above analysis (see the Method section details) with scrubbing different amounts of time points (**Fig. S8** and **Fig. S9**), skipping the temporal filtering step (**Fig. S10**), the FIX-ICA procedure (**Fig. S11**), both the temporal filtering step and the FIX-ICA procedure (**Fig. S12**), or the GSR on the computation of functional connectivity (**Fig. S5**), and convolving FD (**Fig. S16**) or DVARS (**Fig. S13**) with a hemodynamic response function, and the major result, i.e., the motion-based scrubbing effect diminished with controlling the DI effect, remains valid. The finding suggests that the effect of motion-based scrubbing on rsfMRI connectivity should be attributed to time points of high DI values, which are closely related to arousal modulations.

**Figure 4.**
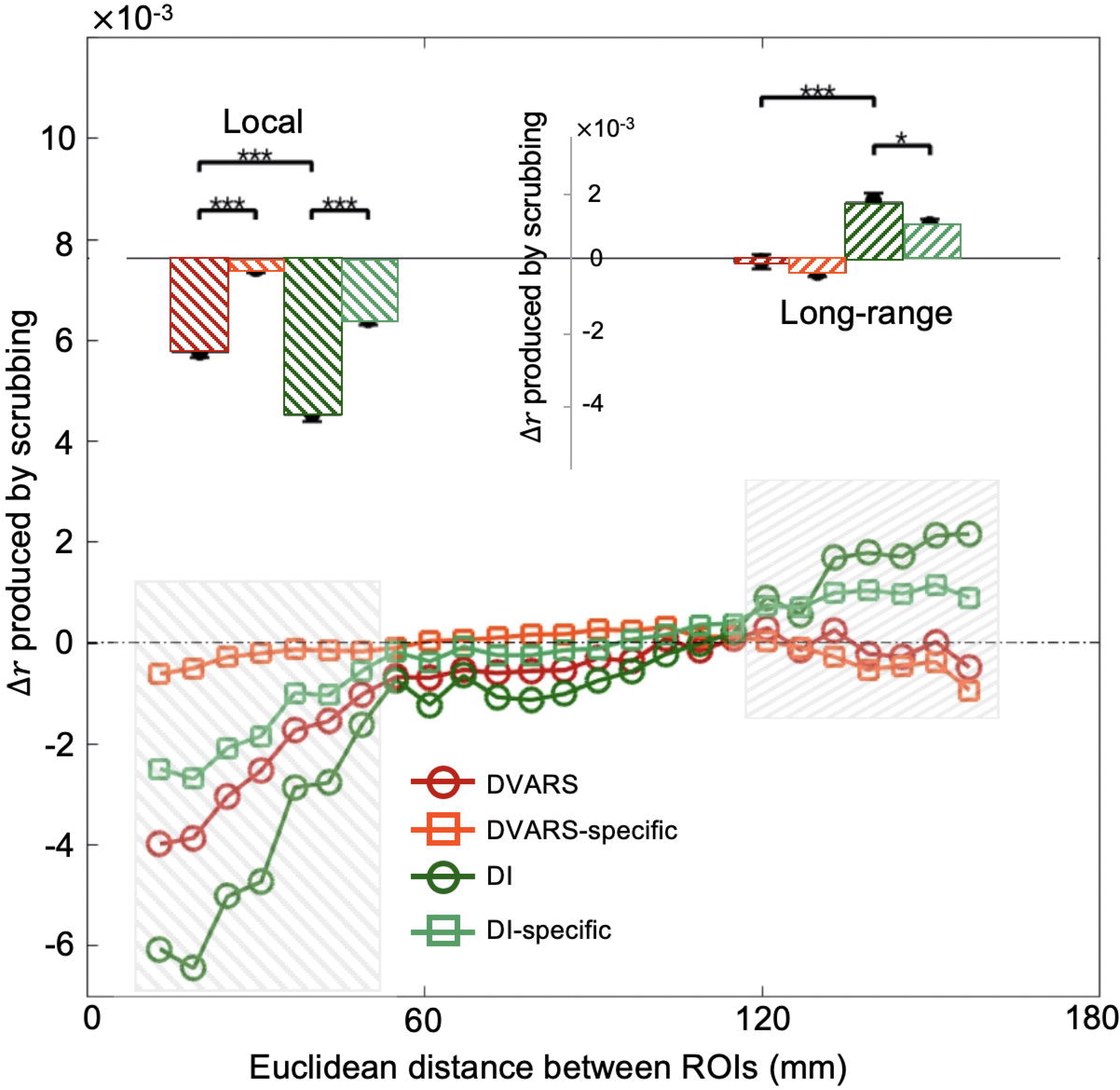
The effect of the DVARS-based scrubbing on rsfMRI connectivity diminished with retaining the high DI volumes. After removing the overlapped time points (9.81% out of 25%) between the DVARS and DI masks from temporal scrubbing, i.e., retaining them for assessing rsfMRI connectivity, the DVARS-specific scrubbing (orange square) produced marginal effects on the local (*t* = 3.19; *p* = 0.00746; *d* = 0.21; df = 468) and long-range (*t* = 3.73; *p* = 0.001339; *d* = 0.24; df = 468) rsfMRI connectivity compared to control group. In contrast, the effect of the DI-specific scrubbing (light green square) remains significant on either local (*t* = 12.80; *p* = 0; *d* = 0.83; df = 468) or long-range (*t* = 5.86; *p* = 0; *d* = 0.38; df = 468) rsfMRI connectivity compared to control group. The colored circles/squares represent the averaged connectivity changes every 6 mm. The local (ROI pairs with a distance between 13 and 49 mm, gray box) and long-range (ROI pairs with a distance between 125 and 161 mm, gray box) rsfMRI connectivity changes are summarized as a bar plot and shown as an inset in the top region. The error bar represents SEM across subjects. The asterisks represent the level of significance: *, 0.01 < *p* ≤ 0.05, and ***, *p* ≤ 0.001.

Since a recent study suggested the motion-based scrubbing effect should be largely attributed to associated respiratory modulations (Power et al. 2018), we further calculated the respiratory volume (RV) signal (Chang et al. 2009) for a subgroup of HCP subjects (N = 407 out of 469 subjects) with available physiological data, and then estimated its scrubbing effect on rsfMRI connectivity. Similar to the DVARS results, scrubbing the high RV time points results in smaller changes in the local (*t* = 14.02; *p* = 0; *d* = 1.00; df = 406) and long-range (*t* = 6.26; *p* = 2.95×10^−6^; *d* = 0.43; df = 406) rsfMRI connectivity compared with the DI-based scrubbing, and this effect completely disappear with retaining the overlapped time points with the DI mask (**Fig. 5**). We also convolved the RV with a respiratory response function (Birn et al. 2008) before doing the scrubbing analysis and a similar result was obtained (**Fig. S14**).

**Figure 5.**
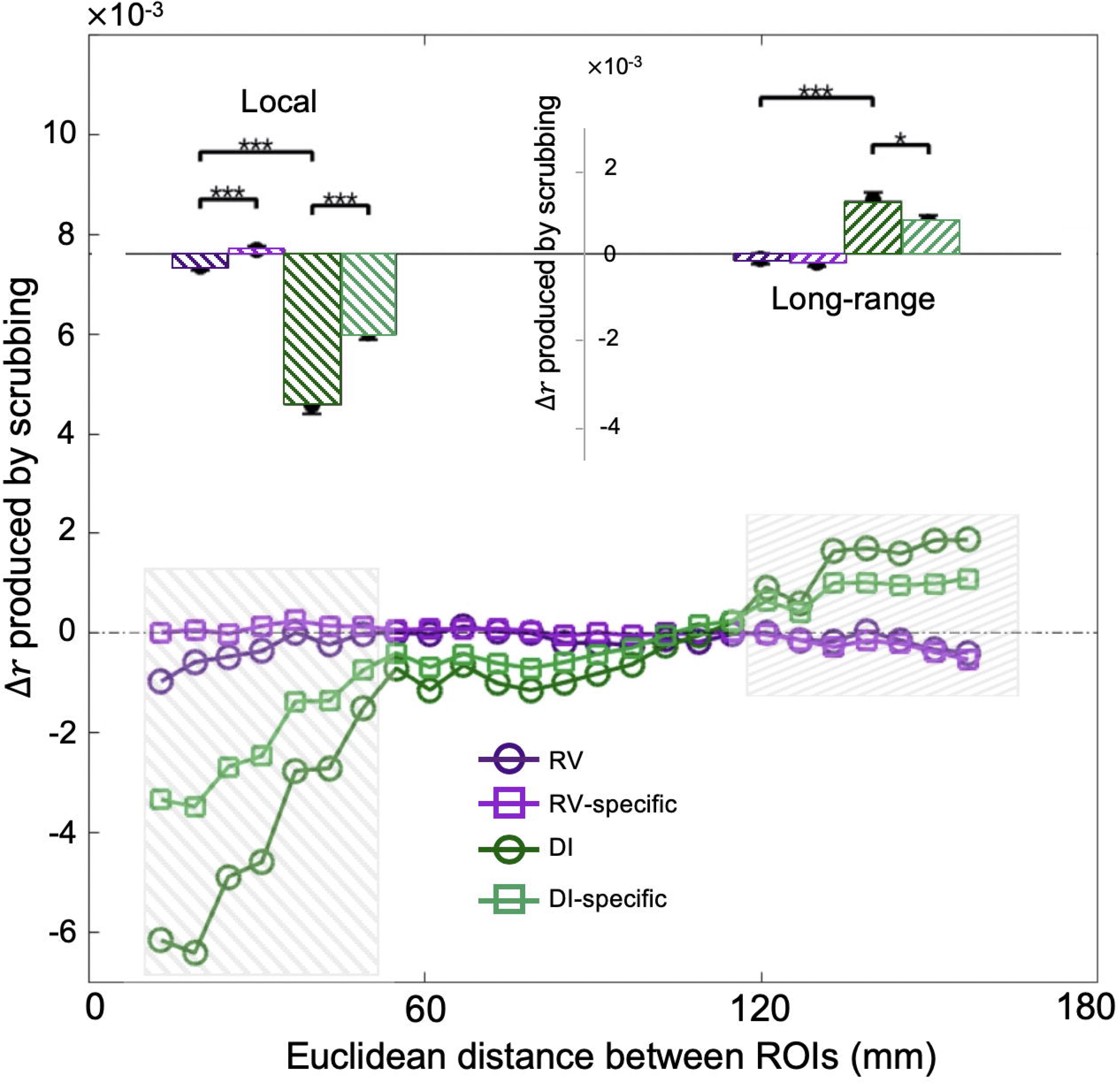
The effect of the RV-based scrubbing on rsfMRI connectivity diminished with retaining the high DI volumes. Note that RV was not convolved with the respiratory response function before the scrubbing analysis. The RV-based scrubbing produced marginal significant effects on local (*t* = 3.34; *p* = 0.001; *d* = 0.23; df = 406) and non-significant effects on long-range (*t* = 1.77; *p* = 0.0766; *d* = 0.12; df = 406) rsfMRI connectivity compared to control group. After removing the overlapped time points (6.84% out of 25%) between the RV and DI masks from temporal scrubbing, i.e., retaining them for assessing rsfMRI connectivity, the effect of the DI-specific scrubbing (light green square) remains significant on either local (*t* = 15.09; *p* = 0; *d* = 0.99; df = 406) or long-range (*t* = 4.61; *p* = 6.41×10^−7^; *d* = 0.31; df = 406) rsfMRI connectivity compared to control group. The colored circles/squares represent the averaged connectivity changes every 6 mm. The local (ROI pairs with a distance between 13 and 49 mm, gray box) and long-range (ROI pairs with a distance between 125 and 161 mm, gray box) rsfMRI connectivity changes are summarized as a bar plot and shown as an inset in the top region. The error bar represents SEM across subjects. The asterisks represent the level of significance: *, 0.01 < *p* ≤ 0.05, and ***, *p* ≤ 0.001. RV, respiratory volume.

### Temporal dynamics of various metrics around large head motions

A detailed temporal relationship among various metrics would help us understand their specific role in affecting rsfMRI connectivity. Cross-correlation functions calculated between all possible pairs of FD, DVARS, and DI revealed a rather complicated temporal relationship among them. The cross-correlation function between DI and DVARS had a single peak (the peak Pearson’s correlation *r* = 0.37, *p* = 0 with df = 467) at the zero delay (**Fig. 6A**, left), whereas FD and DI reached their maximum correlation (the peak Pearson’s correlation *r =* 0.055, *p* = 0.030 with df = 467) with DI being shifted ahead by 9.36 seconds, i.e., the DI is lagging behind FD (**Fig. 6A**, middle). More interestingly, the cross-correlation function for the FD-DVARS pair showed two separate peaks with one at the zero delay (the peak Pearson’s correlation *r* = 0.13, *p* = 4.07×10^−6^ with df = 467) and the other with a significant delay (the peak Pearson’s correlation *r* = 0.10, *p* = 3.13×10^−4^ with df = 467) (**Fig. 6A**, right). The peak correlation between FD and DI is smaller than those of the other two pairs.

**Figure 6.**
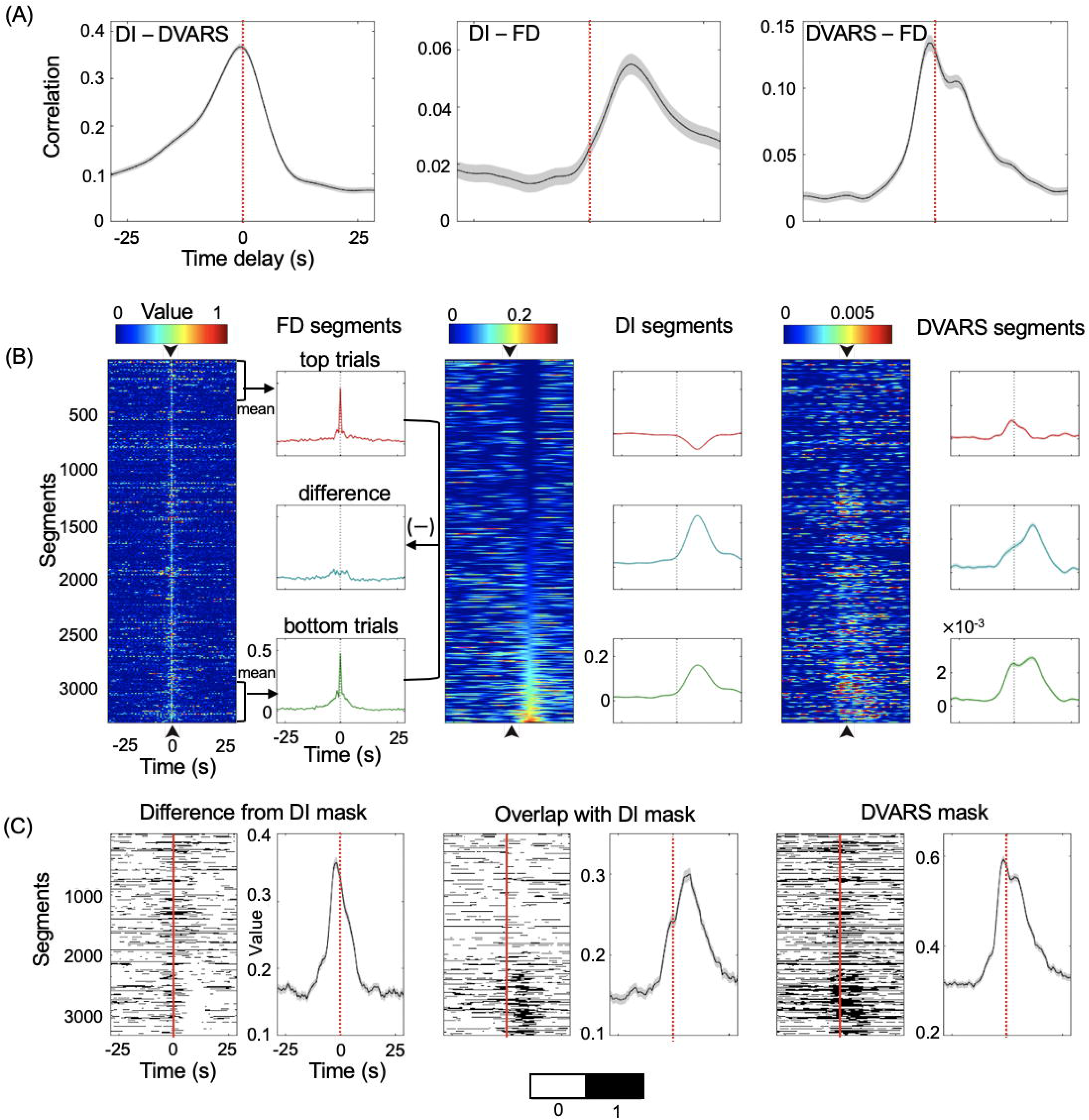
Temporal relationships among FD, DVARS and DI. (A) Cross-correlation functions of FD, DVARS, and DI. A zero-lag peak appears at the DI-DVARS correlation function (left); the maximum correlation between FD and DI appears with 9.36 seconds delay (middle); and the DVARS-FD correlation function shows two distinct peaks (right). The cross-correlation functions are averaged across subjects with the shaded region representing area within 1 SEM. (B) Segments of FD (left), DI (middle), and DVARS (right) were extracted with respect to the 3331 identified FD spikes (time 0) and sorted according to the DI values at 9.36 seconds. The top (red) and bottom (green) 400 segments were averaged and their differences (cyan) were also calculated. The shadow represents regions within one SEM. The black arrows and dash lines indicate time zero. (C) Corresponding segments of the DVARS-based mask (right), its overlap with (middle) and difference from (left) the DI-based mask was also extracted and shown. The average of these segments was also shown on the right side of each panel with the shadow representing regions within one SEM. The red lines indicate time zero.

In order to understand this complex relationship, we employed a trial-based analysis to examine the relative timing of large changes in various metrics, as well as the variability of this relationship over time. The feasibility of this analysis lies in the observation that FD often shows large, abrupt spikes that appear to drive its correlation with DVARS and DI (**Fig. S17**). We identified 3331 FD spikes by finding FD peak points that are 3 standard deviations above the mean. Removing these spikes (4.2 % of the total points) would bring down the covariance between FD and DVARS by 79 % (from 0.0169 to 0.0036). We then extracted FD, DVARS, and DI segments centering on the FD spikes and sorted these segments according to DI values at 9.36 seconds, where we expected to see large peaks according to the FD-DI correlation function (**Fig. 6A**, middle). We found that only a subset of the FD spikes is followed by a DI peak with a delay of 9.36 seconds (**Fig. 6B**, bottom trials). For this group of trials, the DVARS show two separate peaks that are corresponding to the FD spike and the following DI peak respectively (**Fig. 6B**, right panel, bottom trials). In comparison, there are FD spikes of the same amplitude that are associated with only a zero-lag DVARS peak but not the delay DI peak (**Fig. 6B**, top trials). In fact, a majority of FD spikes (~75%) are not associated with any obvious modulations in either DI or GSA (**Fig. 6B** and **Fig. S18**). We further displayed the DVARS scrubbing mask (**Fig. 6C**, right), as well as its overlap with (**Fig. 6C**, middle) and difference from (**Fig. 6C**, left) the DI scrubbing mask for all the trials. As expected, the overlap between the DVARS and DI masks (**Fig. 6C**, middle), which is responsible for the scrubbing effect of DVARS on rsfMRI connectivity (**Fig. 4**), corresponds mostly to the delayed DI/DVARS peaks following only a subset of FD spikes (**Fig. 6B**, bottom trials). In other words, the head motions that are not followed by arousal-related fMRI changes, i.e., the large DI peaks, do not cause significant changes in rsfMRI connectivity (top trials in **Fig. 6B** and **6C**). The above analysis was repeated with a second version of DI that was calculated with excluding the entire sensorimotor cortex, and the major results remained the same (**Fig. S19**), suggesting that the DI peaks following the subset of FD spikes (bottom trials in **Fig. 6B** and **6C**) are unlikely caused by sensorimotor activations related to the head motion. Instead, the FD spikes and associated DI peaks in this subset of trials likely originate from transient arousal modulations, and their relative delay may reflect the hemodynamic delay between neural/behavioral signals and fMRI, as well as their relative phases in the transient arousal event (~10-15 seconds) (Liu et al. 2018). The RV segments were also extracted around the FD spikes (**Fig. S20**). A large and sharp RV modulation was observed in phase with the FD spikes associated with the DI peaks (bottom trials in **Fig. 6B**) but not present in those sole FD spikes (top trials in **Fig. 6B**), suggesting the co-occurrence of respiration modulations with both head motion and arousal modulation.

## Discussion

Here we showed that the association between the head motion parameters and rsfMRI connectivity, which has been interpreted as a causal relationship (Power et al. 2012, 2014, 2018; Satterthwaite et al. 2012; van Dijk et al. 2012), may arise spuriously from transient arousal modulations. We first developed an fMRI-based arousal index, i.e., DI, to track arousal modulations and validated this index through its correlations with arousal-related behavioral measures. We then showed that the effect of temporal scrubbing based on the motion parameters, i.e., FD and DVARS, on rsfMRI connectivity is actually attributed to time points associated with transient arousal modulations. Finally, we further elucidated complex temporal relationships among the motion and arousal indices using a trial-based analysis.

The association between the head motion parameters, i.e., the FD and DVARS, and rsfMRI connectivity may arise from arousal modulations in two ways. First, a significant proportion of this relationship comes spuriously from the use of DVARS, a widely used index for quantifying head motion based on fMRI signal itself. The DVARS index is defined as the root mean square of whole brain fMRI signal changes between consecutive time points and thus sensitive to any large, widespread fMRI changes (Smyser et al. 2010). Although it was designed to detect head motion, it would also be sensitive to transient arousal modulations that have been shown to induce large, global fMRI changes (Fukunaga et al., 2006; He and Liu, 2012; Kiviniemi et al., 2005; Licata et al., 2013; Wong et al., 2016, 2013). The two DVARS peaks that correspond to the FD spike and following DI peak in our trial-based analysis (**Fig. 6B**, bottom trials) likely represent these two types of DVARS modulations. On the other hand, the observed coupling of the FD spikes and DI peaks (**Fig. S17**), though with a significant time delay, could be the second source of the motion-rsfMRI relationship. Only a small proportion of DI peaks were associated with FD spikes (**Fig. 6B**, bottom trials), which might account for the marginal scrubbing effect based on FD (**Fig. 2**), derived from image alignment parameters and thus closely related to real head motions. Another major reason for the relatively weak effect of the FD-based scrubbing is likely the significant time delay between the FD spikes and associated DI peaks (**Fig. 6B**, bottom trials), since convolving FD with a hemodynamic response function significantly improved its scrubbing effect to a level close to the DVARS-based scrubbing (**Fig. S16**). It is worth noting that the previous study (Power et al. 2012) has augmented their FD/DVARS scrubbing masks temporally by 1 frame back and 2 forward. The forward augmentation by two frames (~5 seconds) is expected to achieve a similar effect as the convolution. The sleepiness induced by sleep deprivation can lead to serious head motions (Van Den Berg 2006). Similar to the coupling between the FD and DI peaks, this relationship could be caused by transient arousal modulations and associated physiological changes during the drowsy and sleepy states. It is also worth noting that the DI peak following the FD spike should not be attributed to motion-related sensorimotor activity and associated fMRI changes since removing the entire sensorimotor cortex from the DI calculation had almost no effects on the result (**Fig. S19**).

The contribution of arousal modulations to the spurious relationship between rsfMRI connectivity and head motion parameters explains a few puzzling observations. First, the effect of head motion on rsfMRI connectivity persists for, and even reach its peak at, more than 10 seconds (Power et al. 2014; Byrge and Kennedy 2018). Although this observation has been explained from the perspective of spin-history artifacts, the explanation may not be satisfactory because the spin-history artifact induced by a brief motion should not last for so long and show its peak effect after 10 seconds (Yancey et al. 2011). This prolonged effect is even longer than a typical hemodynamic delay between neural and fMRI signals. However, this characteristic time is roughly consistent with the delay (~9.36 seconds) we observed here between FD spikes and DI peaks. The co-occurrence of FD spikes and large fMRI changes (the large DI) could be mediated by transient arousal events (Liu et al. 2018), and their relative time delay may include both the hemodynamic delay (5~6 seconds) and their relative phases in the arousal event (on the scale of seconds) (Liu et al. 2018). Secondly, the motion-related rsfMRI connectivity changes are not only distance-dependent but also show very systematic spatial patterns (Power et al. 2012, 2018; Yan et al. 2013). Although this is hard to understand from the perspective of head motion, which is relatively random, it is consistent with the sensory-dominant fMRI co-activations associated with arousal modulations, which are expected to promote the local connectivity at the sensory/motor regions.

Transient arousal modulations may also mediate the correlations between rsfMRI and physiological signals, i.e., the heart rate and respiratory volume, which have been widely observed and regarded as evidence for non-neuronal contributions to rsfMRI (Birn et al. 2006; Chang et al. 2013; Özbay et al. 2018). In fact, the recent study (Power et al. 2018) has attributed the motion effect on rsfMRI connectivity to motion-associated respiratory changes, suggesting the motion-rsfMRI relationship and physiology-motion relationship could be unified to a great extent. Consistent with this notion, we observed the co-modulation of rsfMRI signals, head motion, and RV in a subset of data segments responsible for the scrubbing effect (bottom trials in **Fig. 6B** and **Fig. S20**). The physiology-rsfMRI correlation is significant only in a sleep-conducive eyes-closed state but not under a more alert eyes-open condition, and also dependent on the EEG alpha power, an indicator of brain vigilance state (Yuan et al. 2013). The DI modulations following large RV peaks (**Fig. S20**) suggested a sensory-dominant fMRI co-activation pattern with very specific de-activations at the DMT and the NB, the subcortical regions known to get involved in arousal regulation (Buzsaki et al. 1988; Saper et al. 2005; Posner, J., Saper, C., Schiff, N., & Plum 2008; Komlosi 2015). It is worth noting that a similar sensory-dominant pattern is also present in rsfMRI correlations to cardiac signals (Özbay et al. 2018), as well as in electrocochleography (ECoG) gamma-power changes at global rsfMRI peaks (Liu et al. 2015). Thus, the co-modulation of rsfMRI and RV signals might be driven by, or at least associated with, the transient arousal modulations. In turn, the head motion might be the secondary effect caused directly by the large respiratory modulations, i.e., deep breaths. However, these interpretations remain speculative before proven. Recent studies (Tong and Frederick 2012; Posse et al. 2013; Kiviniemi et al. 2016; Fultz et al. 2019) have reported large rsfMRI changes associated with brain pulsations of different time scales, particularly those infra-slow ones on the timescale of tens of seconds. Such infra-slow brain pulsations are also brain-state dependent and much stronger during sleep, and they are even associated with changes in cerebrospinal fluid (CSF) flow (Fultz et al. 2019). The relationship between the arousal-related rsfMRI and physiological changes and such infra-slow brain pulsations is unclear and remains a challenge for future research.

One should also be cautious about the confounding effects of the arousal in rsfMRI studies of various brain diseases, a significant portion of which are either associated with disrupted sleep and circadian rhythms or treated with medicines that can modulate brain arousal state (Whitfield-Gabrieli et al. 2009; Breen et al. 2014; Cohen et al. 2014; Yang et al. 2014; Musiek et al. 2015). It is important to differentiate rsfMRI changes caused by disease-related brain reorganization from those merely reflecting disease-associated vigilance changes. Arousal may affect cognitive performance given its known role in attention regulation and information processing (Pribram and McGuiness 1975), and the ability to regulate arousal state may also be correlated with other subject traits (Lagarde and Batejat 1994). Therefore, transient arousal modulations might be partly responsible for the correlation observed between rsfMRI connectivity/dynamics and behavioral measures (Smith et al. 2015; Vidaurre et al. 2017). Given these potential confounding effects of the arousal, researchers should be cautious about large global fMRI signals, high DI values, and the sensory-dominant pattern in their rsfMRI findings, which are all indicators of arousal-related fMRI changes.

Properly controlling the arousal effect could be important for using rsfMRI to measure functional brain connectivity. Prospective studies need to better control and monitor arousal fluctuations during resting-state scans, which can be achieved by, for example, avoiding rsfMRI experiments under the sleep-conducive eyes-closed condition and/or including EEG or eyelid video with fMRI acquisitions. Retrospective analyses on existing datasets should try to minimize the arousal’s influence with appropriate post-processing procedures. Although the global signal regression can largely suppress the arousal-related component (Falahpour, Nalci, et al. 2018), its residual effect on rsfMRI connectivity may still remain (Gotts et al. 2013), presumably due to its non-uniform contributions across brain regions, i.e., the sensory-dominant pattern. Given that the arousal modulations take the form of transient events that are temporally separable, removing arousal-contaminated time points from the analysis can be an option for reducing the arousal’s influence. The DI used in this study and other fMRI-based arousal indices (Chang et al. 2016; Falahpour et al. 2018) are preferred metrics, as compared with the motion metrics, for such temporal censoring procedures and expected to locate arousal-related fMRI changes more accurately since they were designed for this purpose. They are also expected to outperform the GSA with incorporating the spatial information of arousal-related fMRI changes, and this is supported by the larger scrubbing effect of DI compared with GSA (**Fig. S21**). On the other hand, even assuming the temporal censoring can effectively remove the direct effect of the transient arousal modulations, it is unclear whether the brain connectivity/dynamics are also modulated outside of the time periods of those arousal events under drowsy and sleepy conditions. A more conservative way of controlling the arousal’s effect would be to completely discard sessions/subjects showing large arousal modulations, which can be assessed again using the fMRI-based arousal indices, such as DI, if no other arousal measures available. Alternatively, this could be done by including individuals’ arousal score as a nuisance variable into the statistical model that compares rsfMRI metrics under different groups or brain conditions.

The DI was used to locate arousal-related fMRI changes in this study. It was derived by correlating individual fMRI volumes with the global co-activation pattern induced by an electrophysiological event of transient arousal modulation (Liu et al. 2018). The similar template-matching methods were used for arousal estimation with different templates, including the fMRI correlation maps of the EEG arousal index or behavioral arousal index defined by eyelid positions (Chang et al. 2016; Falahpour, Chang, et al. 2018). These different templates show very similar sensory-dominant patterns and may thus originate from the same event of arousal modulations (Liu et al. 2018). These fMRI-based arousal indices are able to capture both the intra-subject dynamics and inter-subject variation of the brain arousal. They thus can provide a flexible and convenient estimate of the brain arousal to any data sets with rsfMRI data, including recent large-scale imaging initiatives collected from healthy subjects and patients, e.g., the UK Biobank (Miller et al. 2016) and Alzheimer’s Disease Neuroimaging Initiative (ADNI) (Jack et al. 2008). Together with other modalities of data, this may create new opportunities for understanding the mechanisms of arousal regulation as well as its role in the development of many neurological diseases of arousal relevance, e.g., Alzheimer’s disease (Carvalho et al. 2018; Musiek et al. 2018).

There are some potential limitations in this study. First, the DI was computed from rsfMRI signals since there are no other independent arousal measures available in the HCP data. This may raise a potential concern about the circularity of our results since the DI-based scrubbing effect on rsfMRI connectivity is well expected based on the pattern of the template for DI calculation. However, it should be clarified that the major point of this study is not the DI-based scrubbing effect on rsfMRI connectivity but that the motion-associated rsfMRI connectivity changes are largely due to these arousal-related rsfMRI changes quantified by the DI. Secondly, the study on the fMRI-based arousal measures is still at the early stage (Chang et al. 2016; Falahpour et al. 2018), and more research is needed to further elucidate their relationship with the brain arousal level. Since the DI was designed to detect the events of transient arousal modulations, it likely quantifies the level of drowsiness that reaches the peak at states of intermediate arousal level, such as drowsy and light sleep stages. Therefore, one needs to be cautious when using these fMRI-based metrics to estimate the arousal levels. At the same time, we also want to emphasize that the spatial co-activation pattern that we used for DI calculation has been linked to transient arousal modulation events, the template-based arousal measure using rsfMRI has been validated with simultaneously electrophysiological and behavioral measures (Chang et al. 2016; Falahpour et al. 2018), and we also showed that the DI is much more closely linked to arousal-related behavioral measures, particularly as compared with the motion metrics (**Fig. 2**). Thirdly, it remains unclear whether the head motion of arousal irrelevance can induce systematic changes in rsfMRI connectivity. The scrubbing based on FD, presumably a better parameter for real motion compared with DVARS, produced only weak effects on rsfMRI connectivity (**Fig. S15**), which is likely due to the time delay between the motion and associated fMRI changes (**Fig. 6B**). The exclusion of the high DI time points from scrubbing completely removed this effect (**Fig. S15** & **S16**), suggesting that the pure head motion of arousal irrelevance has limited effects on rsfMRI connectivity. However, it remains unclear whether this would be true for rsfMRI data with more serious head motions, such as those from adolescence and children (Power et al. 2012). Moreover, since we only examined the distance-dependent rsfMRI connectivity modulation, we cannot exclude the possibility that the pure head motion may have very specific effects on individual connections, which were then cancelled out while we averaged across connections of similar distances. Therefore, it remains a challenging issue for future studies to determine whether the head motion irrelevant to arousal can induce systematic changes in rsfMRI connectivity.

In summary, the rsfMRI connectivity changes associated with head motion parameters are not caused by the motion itself but by transient arousal modulations, which have been shown to induce profound fMRI changes. The arousal-related fMRI change may potentially confound the relationships between rsfMRI connectivity and other measures. Caution should be exercised if large global rsfMRI signals, higher values in fMRI-based arousal indices, or sensory-dominant brain maps are found in rsfMRI analyses.

## Contributions

Y.G. and X.L. designed the study and performed the analyses; Y.G., X.L., F.H., and L.S. wrote the paper.

## Funding

This research was supported by the NIH Pathway to Independence Award (K99/R00) 4R00NS092996-02.

## Acknowledgements

The authors declare no competing financial interests and we have no conflict of interest to declare. We thank Dr. Matthew F. Glasser for sharing the information about subjects’ sleeping state inside the scanner and Yikang Liu for reviewing the manuscript. We also thank Maryam Falahpour for providing the map of fMRI correlations to EEG vigilance index.

