## Supplemental Figures for "Transient arousal modulations contribute to resting-state functional connectivity changes associated with head motion parameters"

### Supporting Information (SI)

### SI Figures

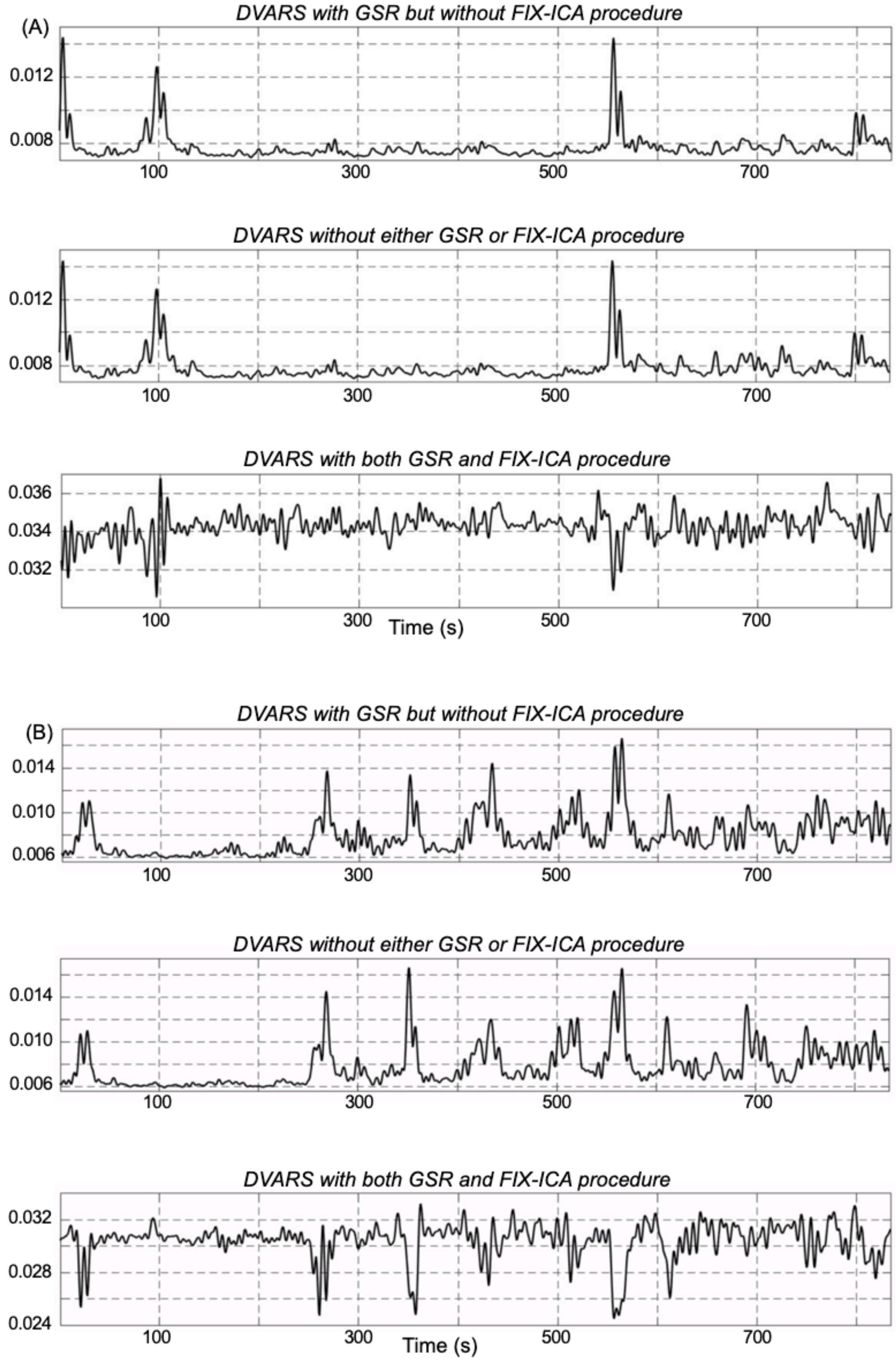

Fig. S1: Different versions of DVARS calculated from two random subjects (A and B). DVARS time courses calculated with applying global signal regression (GSR) but not the FIX-ICA procedure (top panels, A-B) are very similar to those obtained without applying either FIX-ICA procedure or GSR (middle-panels, A-B). DVARS time courses with applying both GSR and FIX-ICA procedure (bottom-panels, A-B), showing sharp dips as indicated in the previous study (Glasser et al. 2018).

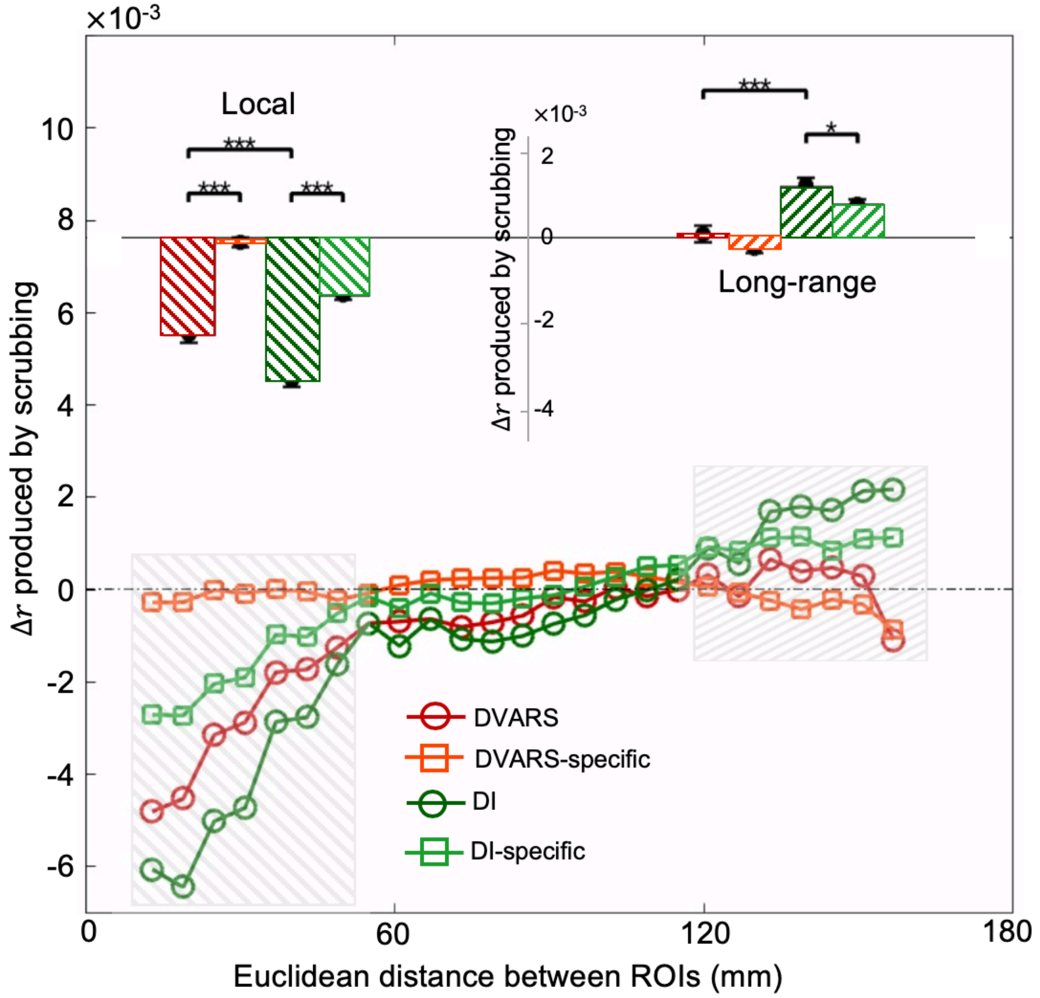


Fig. S2: A similar temporal scrubbing effect of DI and DVARS on rsfMRI connectivity was observed using DVARS calculated without applying global signal regression (GSR) compared with that using DVARS calculated with applying GSR in Fig. 4. The percentage of overlapped time points between the DVARS and DI masks from temporal scrubbing was 10.23% of the total time points. The DVARS-based scrubbing (red circle) produced significant effects on the local (one sample $t = 11.70$; $p = 0$; Cohen's $d = 0.77$; degree of freedom (df) = 468; same as below) and non-significant effects on the long-range ($t = 0.039$; $p = 0.97$; $d = 0.0025$; df = 468) rsfMRI connectivity compared to the control group. The local scrubbing effect of the DVARS-based scrubbing on rsfMRI connectivity diminished with retaining the high DI volumes ($t = 9.89$; $p = 0$; $d = 0.66$; df = 468). These results are consistent with the results from Fig. 4 given that the DVARS calculated without applying GSR is similar to the DVARS with applying GSR (Fig. S1). The colored circles represent the averaged connectivity changes every 6 mm. The local (ROI pairs with a distance between 13 and 49 mm, gray box) and long-range (ROI pairs with a distance between 125 and 161 mm, gray box) rsfMRI connectivity changes are summarized as a bar plot and shown as an inset in the top region. The error bar represents SEM across subjects. The asterisks represent the level of significance: *, $0.01 < p \leq 0.05$ and ***, $p \leq 0.001$.

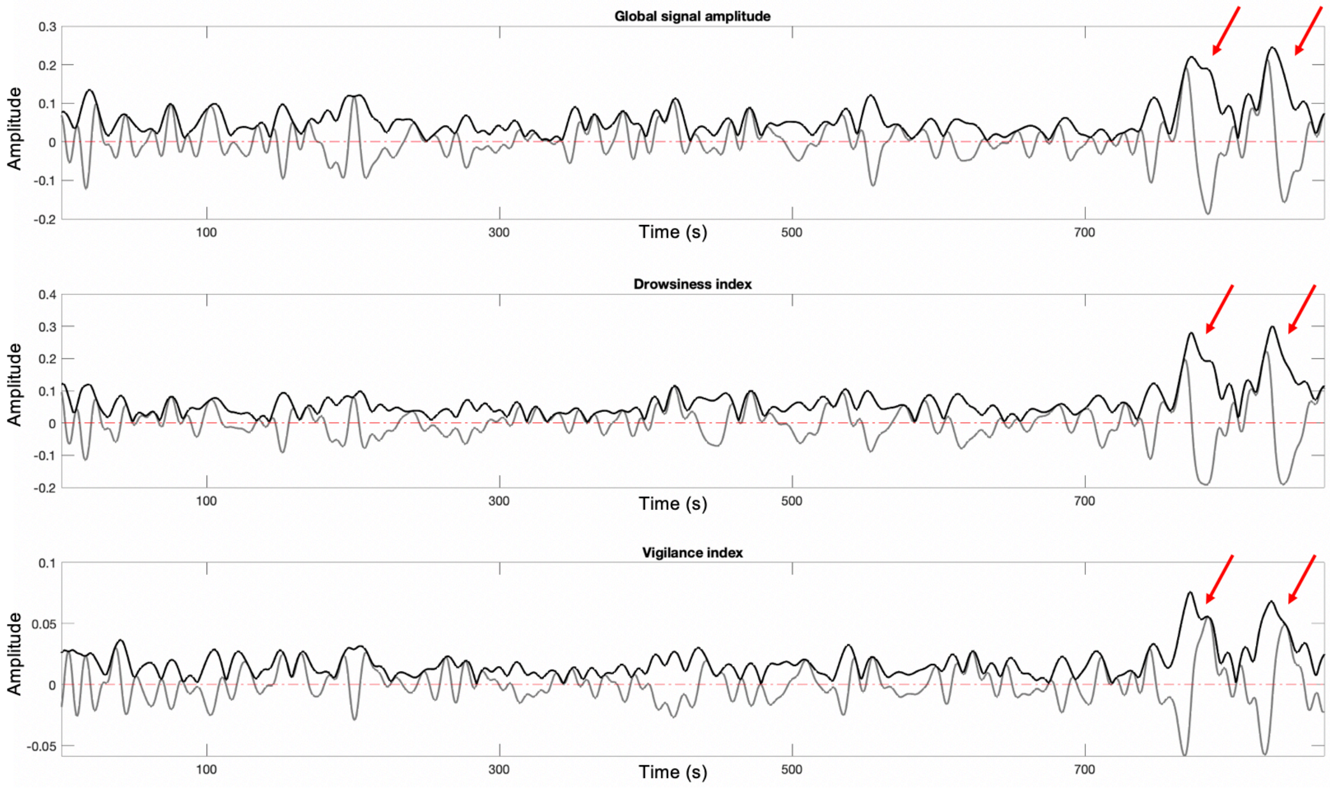


Fig. S3: An example of the global signal amplitude (black, top row), drowsiness index (black, middle row), and vigilance index (black, bottom row) from a subject in the HCP dataset. It can be seen that the large peaks and valleys of the global signal and the correlation to the pre-defined spatial templates (grey lines) often come as pair noted by red arrows. The envelope amplitude captured both peaks and valleys (black, top, middle and bottom rows).

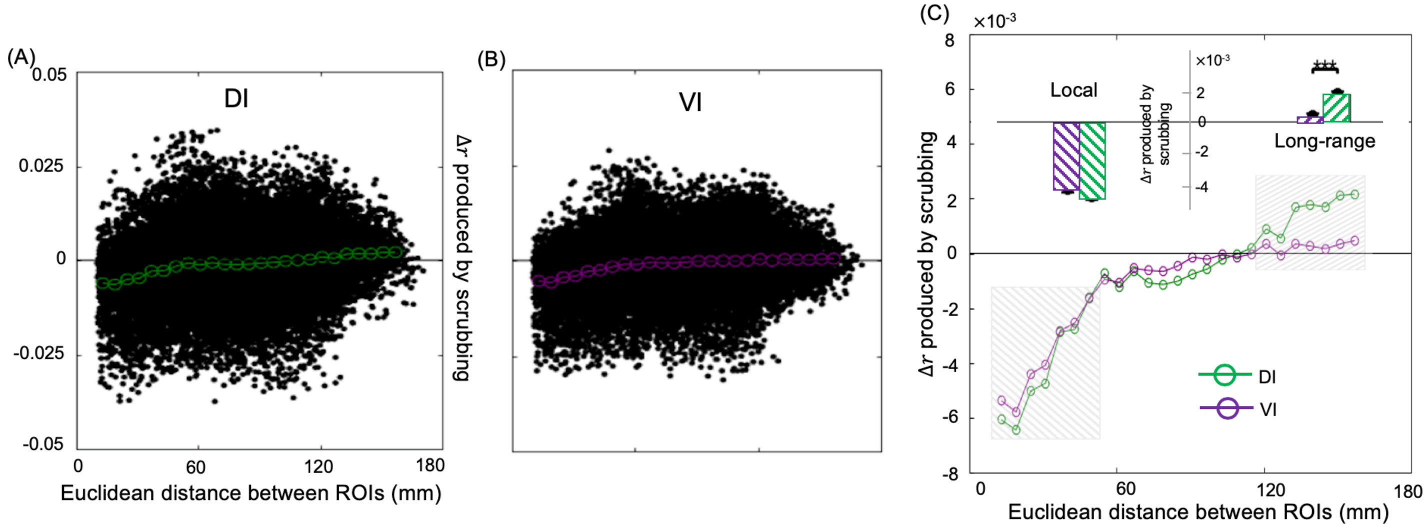


Fig. S4: Applying different spatial templates showed similar scrubbing effects on rsfMRI connectivity. The map of rsfMRI correlations to EEG vigilance index (Falahpour et al. 2018) was used to calculate a vigilance index (VI). Removing time points captured by both drowsiness index (DI) (A) and VI (B) reduced the local but increased the long-range rsfMRI correlations between the 264 pre-defined brain ROIs. A comparison of temporal scrubbing effects (C) suggests that the DI-based scrubbing has a larger effect on the long-range rsfMRI connectivity than VI-based scrubbing ($t = 4.62$; $p = 4.35\text{x}10^{-6}$; $d = 0.31$; df = 468). The colored circles represent the averaged connectivity changes every 6 mm. The local (ROI pairs with a distance between 13 and 49 mm, gray box) and long-range (ROI pairs with a distance between 125 and 161 mm, gray box) rsfMRI connectivity changes are summarized as a bar plot and shown as an inset in the top region (C). The error bar represents SEM across subjects. The asterisks represent the level of significance: ***, $p \leq 0.001$.

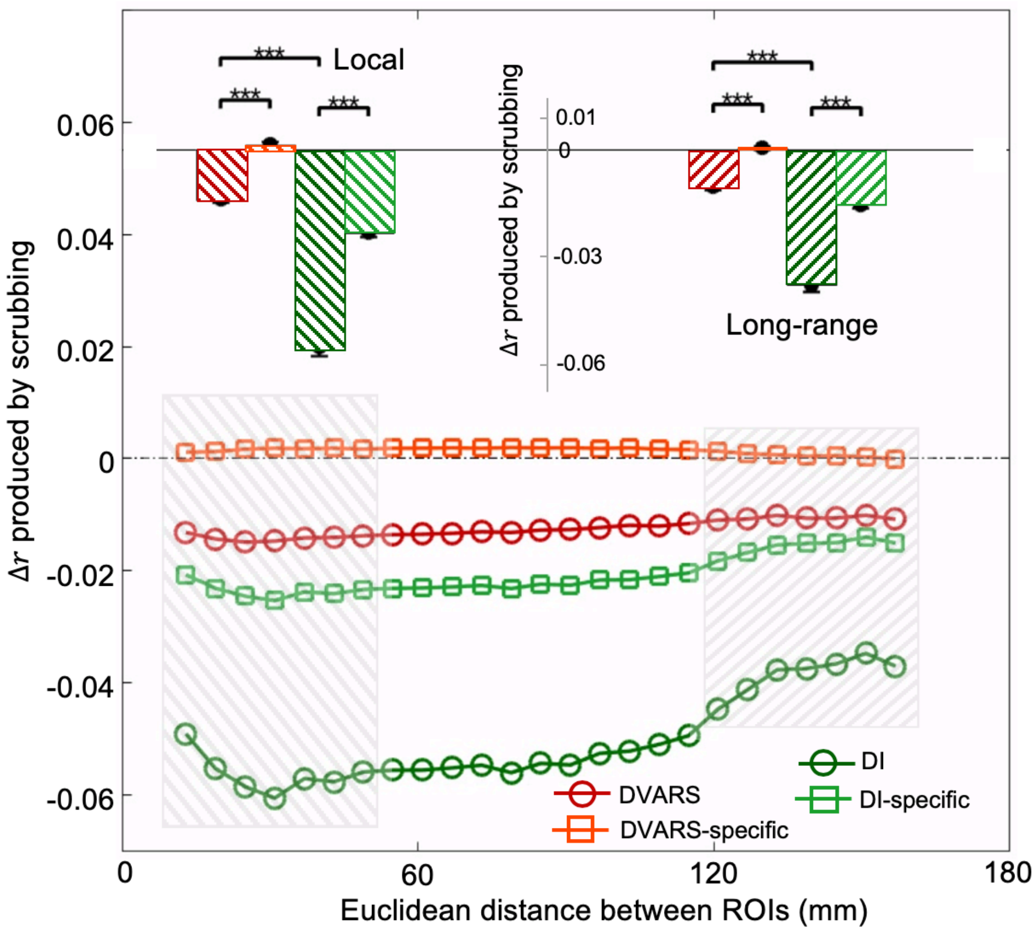


Fig. S5: Compared with the functional connectivity computed by applying the GSR in Fig. 4, here we investigated the scrubbing effect of DVARS and DI using the functional connectivity computed by skipping GSR. The percentage of overlapped time points between the DVARS and DI masks from temporal scrubbing is 9.81% of the total time points. It can be seen that the connectivity changes from the DVARS-based or DI-based scrubbing were not visible and hidden by the global effect. The colored circles represent the averaged connectivity changes every 6 mm. The local (ROI pairs with a distance between 13 and 49 mm, gray box) and long-range (ROI pairs with a distance between 125 and 161 mm, gray box) rsfMRI connectivity changes are summarized as a bar plot and shown as an inset in the top region. The error bar represents SEM across subjects. The asterisks represent the level of significance: ***, $p \leq 0.001$.

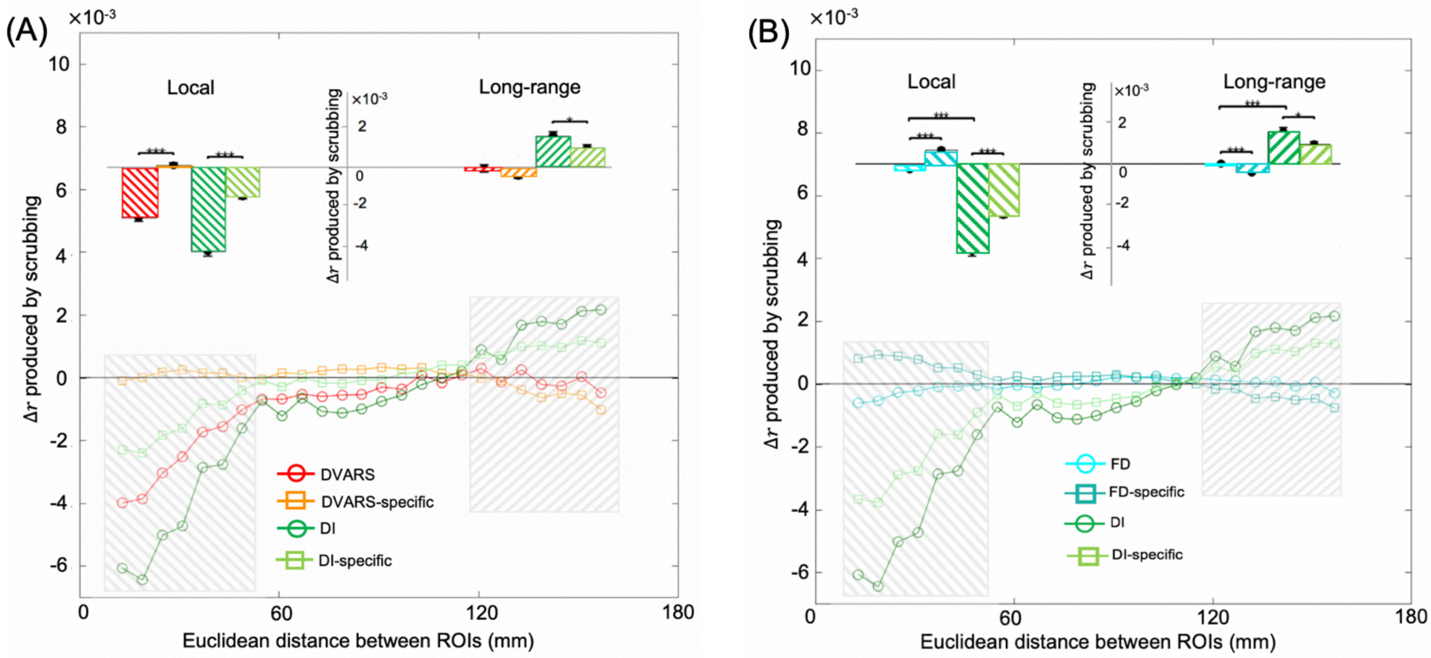


Fig. S6: Two examples using the method to estimate the metric-specific scrubbing effect without controlling the DI effect. The DVARS-specific mask in (A) is calculated as the DVARS mask with excluding its overlap with the DI mask. The same method is applied to calculate the FD-specific mask in (B). The others are the same with the Fig. 4 and Fig. S15 respectively.

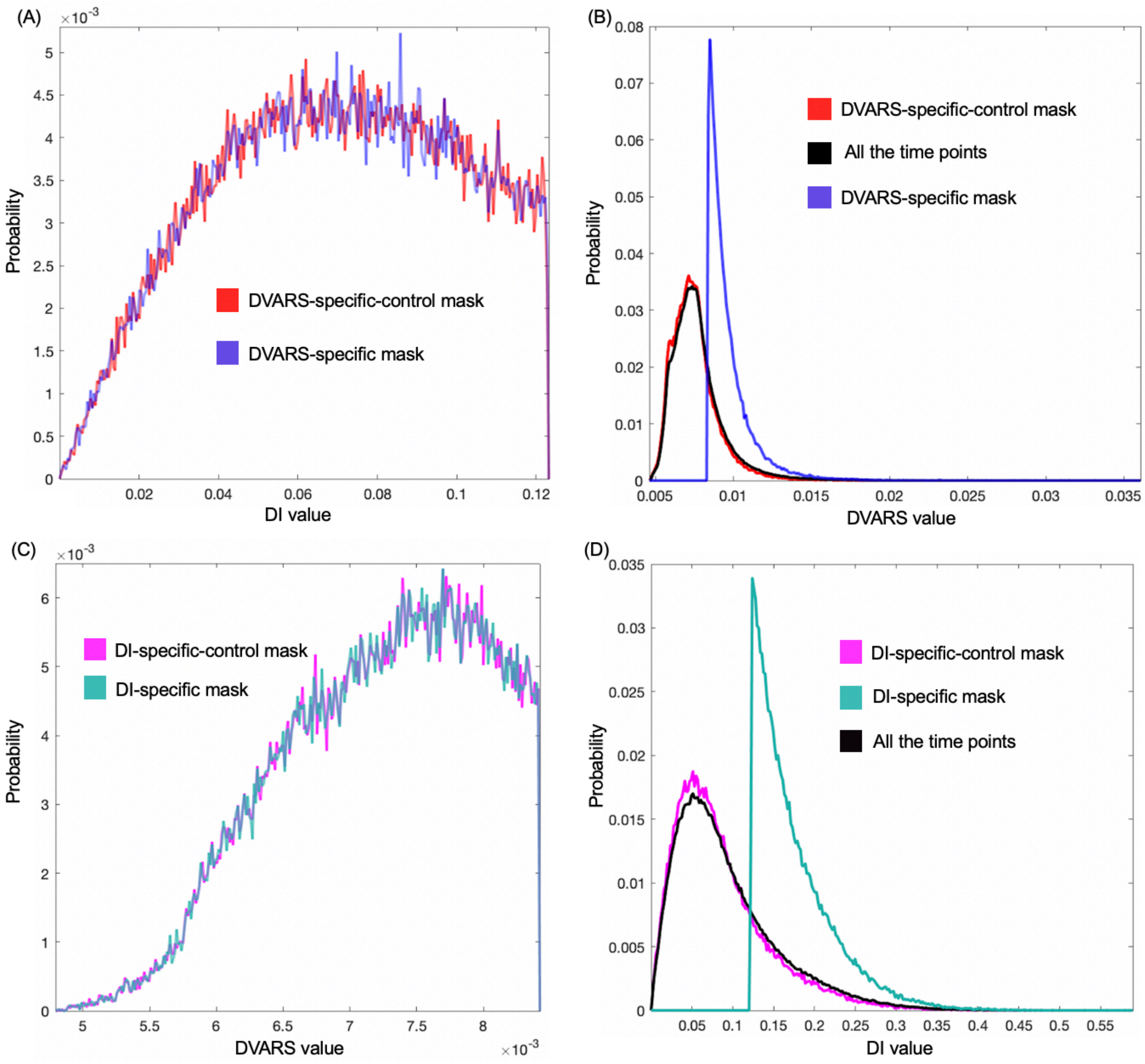


Fig. S7: The DVARS-specific-control and DI-specific-control masks used to estimate the metric-specific scrubbing effect in the Fig. 4. The DVARS-specific-control mask (red, A-B) has a similar DI value with the DVARS-specific mask (blue, A) and a similar DVARS value with all of the time points (black, B). The DI-specific-control mask (magenta, C-D) has a similar DVARS value with the DI-specific mask (dark cyan, C) and a similar DI value with all of the time points (black, D).

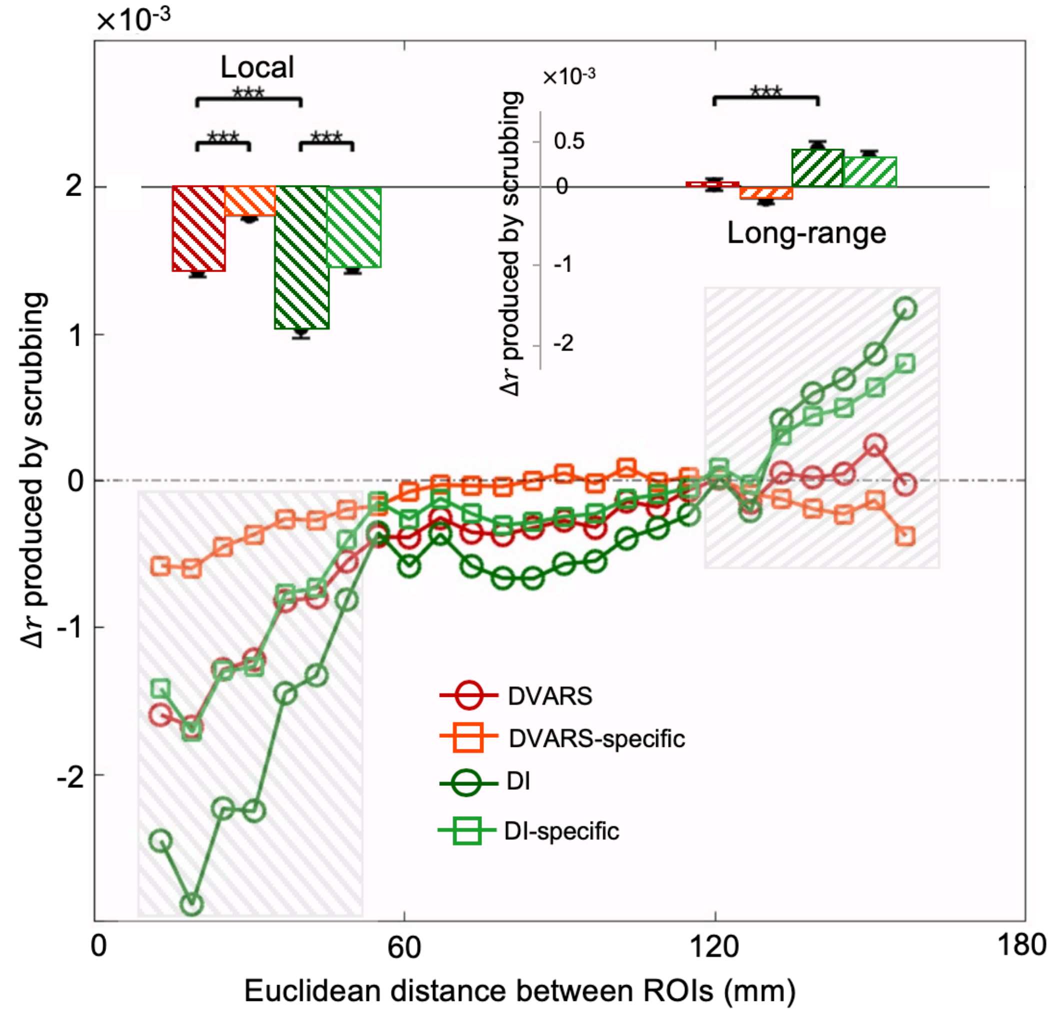


Fig. S8: Compared with the 25% scrubbed time points in Fig. 4, we examined the rsfMRI connectivity changes by scrubbing 10% time points of DVARS or DI. The percentage of overlapped time points between the DVARS and DI masks from temporal scrubbing is 2.3% of the total time points. The DVARS-based scrubbing (red circle) produced significant effects on the local ($t$ = 10.92; $p$ = 0; $d$ = 0.69; df = 468) and non-significant effects on the long-range ($t$ = 0.75; $p$ = 0.45; $d$ = 0.048; df = 468) rsfMRI connectivity compared to control group. The local scrubbing effect of the DVARS-based scrubbing on rsfMRI connectivity diminished with retaining the high DI volumes ($t$ = 7.10; $p$ = 0; $d$ = 0.44; df = 468). The local and long-range functional connectivity changes of scrubbing 10% time points show smaller amplitude than that of 25% time points, given that the amount of scrubbed time points is decreased. The colored circles represent the averaged connectivity changes every 6 mm. The local (ROI pairs with a distance between 13 and 49 mm, gray box) and long-range (ROI pairs with a distance between 125 and 161 mm, gray box) rsfMRI connectivity changes are summarized as a bar plot and shown as an inset in the top region. The error bar represents SEM across subjects. The asterisks represent the level of significance: ***, $p \leq 0.001$.

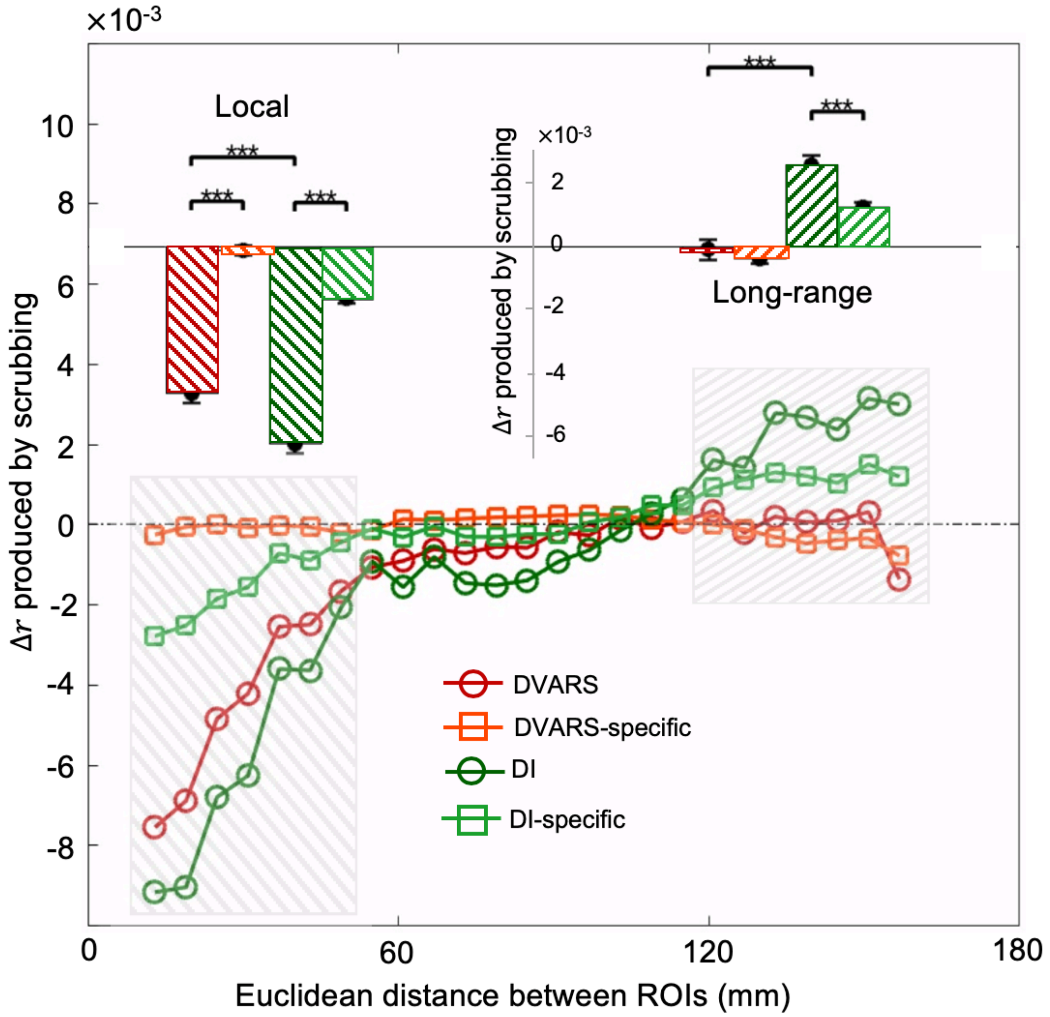


Fig. S9: Compared with the 25% scrubbed time points in Fig. 4, we examined the rsfMRI connectivity changes by scrubbing 40% time points of DVARS or DI. The percentage of overlapped time points between the DVARS and DI masks from temporal scrubbing was 20.12% of the total time points. The DVARS-based scrubbing (red circle) produced significant effects on the local ($t = 15.10$; $p = 0$; $d = 0.98$; df = 468) and non-significant effects on the long-range ($t = 0.64$; $p = 0.52$; $d = 0.04$; df = 468) rsfMRI connectivity compared to control group. The local scrubbing effect of the DVARS-based scrubbing on rsfMRI connectivity diminished with retaining the high DI volumes ($t = 13.36$; $p = 0$; $d = 0.92$; df = 468). The local and long-range functional connectivity changes of scrubbing 40% time points show larger amplitude than that of 25% time points, given that the amount of scrubbed time points is increased. The colored circles represent the averaged connectivity changes every 6 mm. The local (ROI pairs with a distance between 13 and 49 mm, gray box) and long-range (ROI pairs with a distance between 125 and 161 mm, gray box) rsfMRI connectivity changes are summarized as a bar plot and shown as an inset in the top region. The error bar represents SEM across subjects. The asterisks represent the level of significance: ***, $p \leq 0.001$.

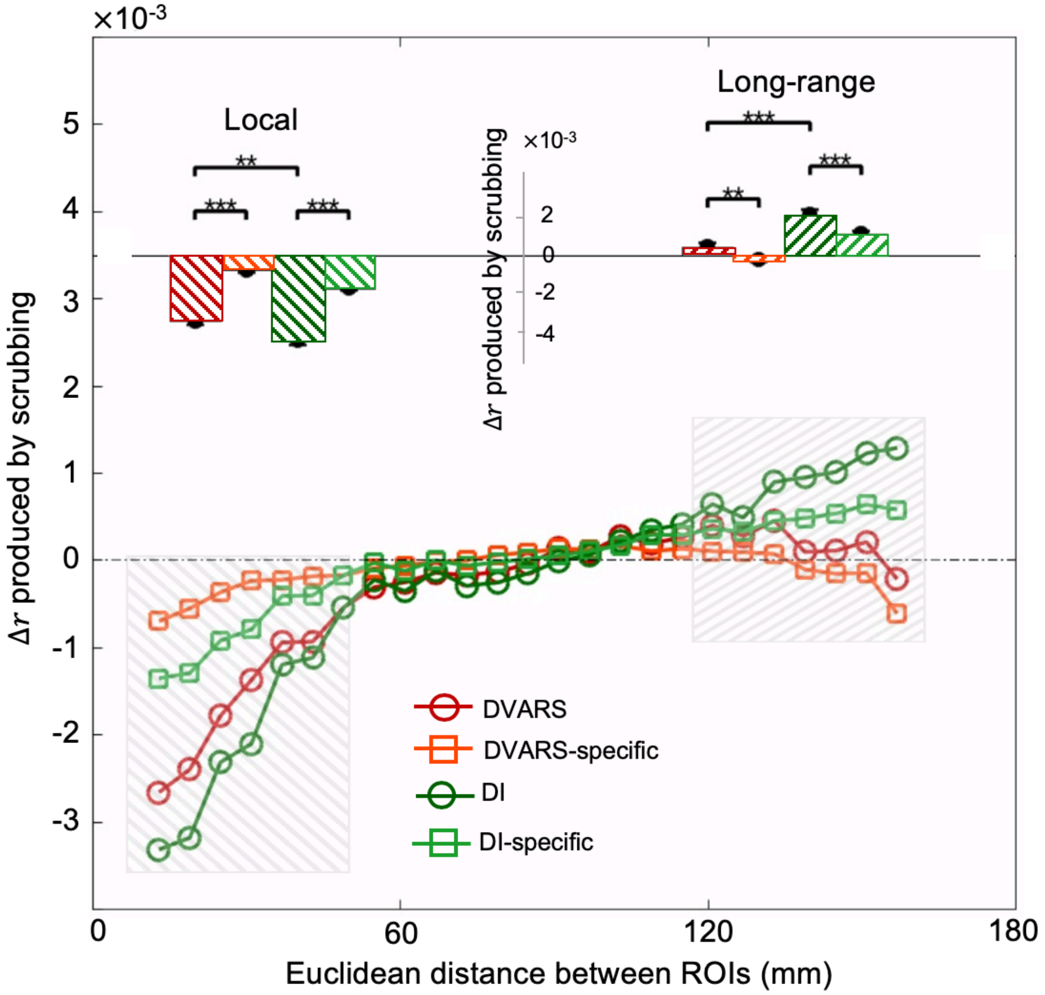


Fig. S10: Compared with the functional connectivity computed using the temporal filter (0.001 - 0.1 Hz) step in Fig. 4, here we investigated the scrubbing effect of DVARS and DI using the functional connectivity computed by skipping the temporal filtering step. The percentage of overlapped time points between the DVARS and DI masks from temporal scrubbing is 9.81% of the total time points. The DVARS-based scrubbing (red circle) produced significant effects on the local ($t$ = 14.08; $p$ = 8.58x10$^{-8}$; $d$ = 0.90; df = 468) and non-significant effects on the long-range ($t$ = 0.68; $p$ = 0.50; $d$ = 0.043; df = 468) rsfMRI connectivity compared to the control group. The local scrubbing effect of the DVARS-based scrubbing on rsfMRI connectivity diminished with retaining the high DI volumes ($t$ = 11.05; $p$ = 0; $d$ = 0.70; df = 468). Compared to the results from Fig. 4, the DI- or DVARS-based scrubbing here generated smaller amplitude of the local and the long-range connectivity changes. We hypothesize that skipping the temporal filtering step might generate smaller connectivity changes of scrubbing. The colored circles represent the averaged connectivity changes every 6 mm. The local (ROI pairs with a distance between 13 and 49 mm, gray box) and long-range (ROI pairs with a distance between 125 and 161 mm, gray box) rsfMRI connectivity changes are summarized as a bar plot and shown as an inset in the top region. The error bar represents SEM across subjects. The asterisks represent the level of significance: **, $0.001 < p \leq 0.01$, and ***, $p \leq 0.001$.

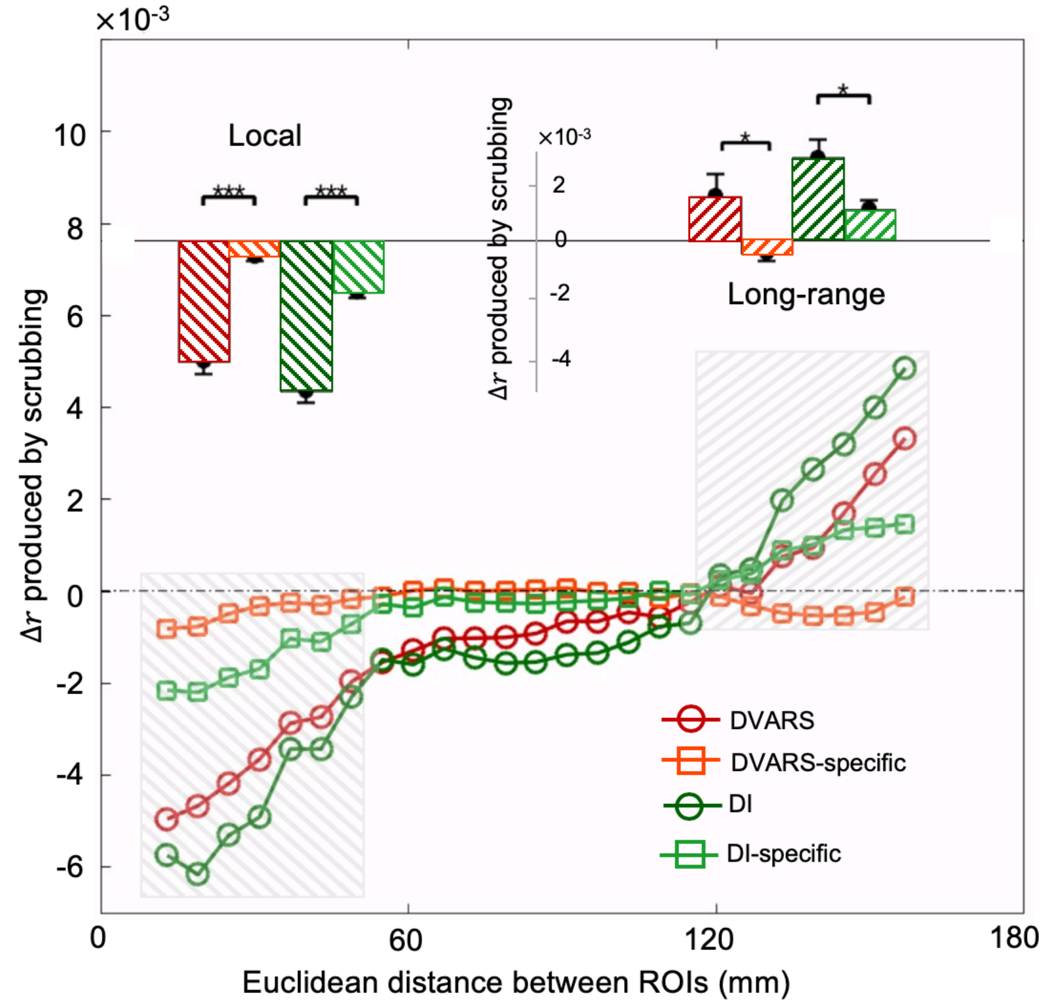


Fig. S11: Compared with the functional connectivity computed using the FIX-ICA procedure in Fig. 4, here we investigated the scrubbing effect of DVARS and DI using the functional connectivity computed by skipping the FIX-ICA procedure. The percentage of overlapped time points between the DVARS and DI masks from temporal scrubbing is 9.81% of the total time points. The DVARS-based scrubbing (red circle) produced significant effects on the local ($t$ = 9.44; $p$ = 0; $d$ = 0.62; df = 468) and non-significant effects on the long-range ($t$ = 1.88; $p$ = 0.061; $d$ = 0.12; df = 468) rsfMRI connectivity compared to control group. The local scrubbing effect of the DVARS-based scrubbing on rsfMRI connectivity diminished with retaining the high DI volumes ($t$ = 7.64; $p$ = 0; $d$ = 0.50; df = 468). Compared to the results from Fig. 4, the DI- or DVARS-based scrubbing here generated larger amplitude of the local and the long-range connectivity changes. We hypothesize that skipping the FIX-ICA procedure might leave the rsfMRI data noisier and therefore, generate larger connectivity changes. The colored circles represent the averaged connectivity changes every 6 mm. The local (ROI pairs with a distance between 13 and 49 mm, gray box) and long-range (ROI pairs with a distance between 125 and 161 mm, gray box) rsfMRI connectivity changes are summarized as a bar plot and shown as an inset in the top region. The error bar represents SEM across subjects. The asterisks represent the level of significance: *, 0.01 < $p \le 0.05$, and ***, $p \le 0.001$.

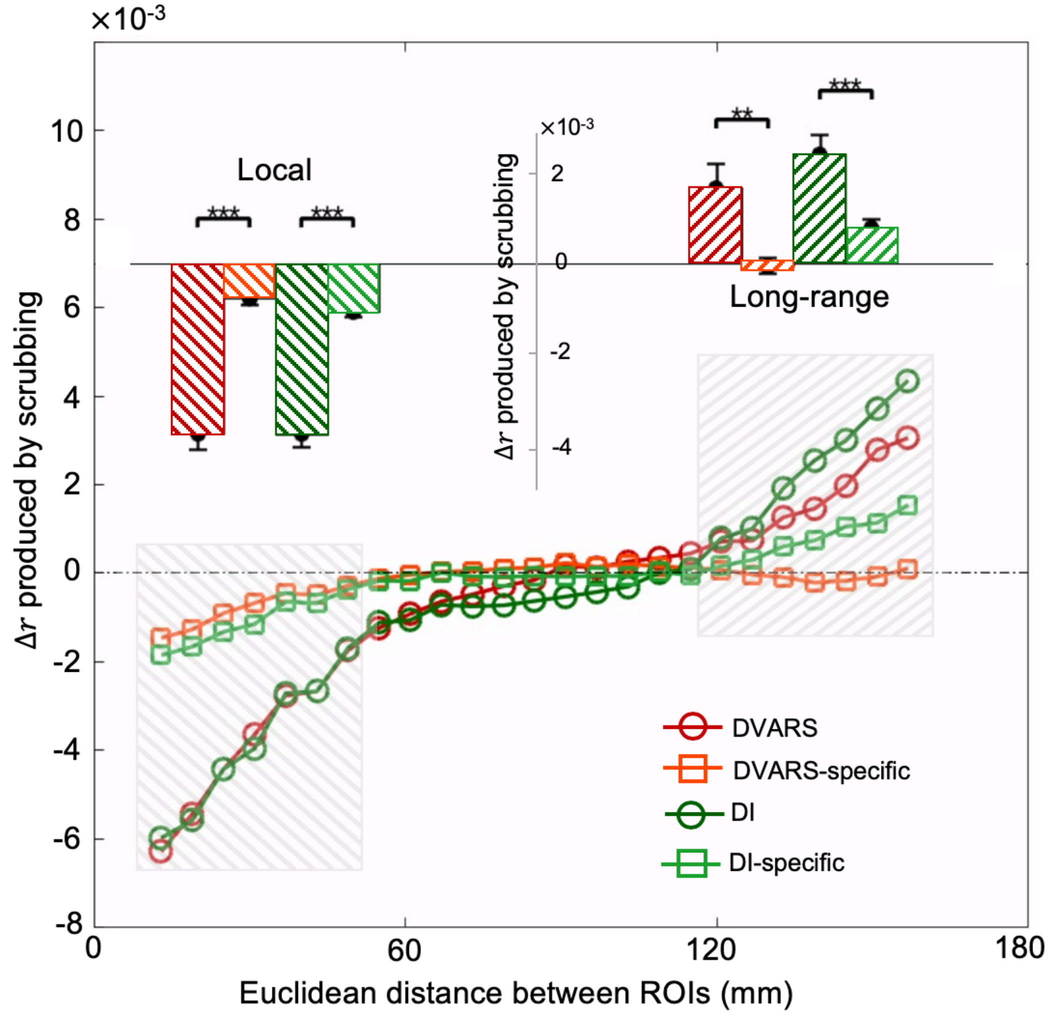


Fig. S12: Compared with the functional connectivity computed using both the temporal filter (0.001 - 0.1 Hz) step and FIX-ICA procedure in Fig. 4, here we investigated the scrubbing effect of DVARS and DI using the functional connectivity computed by skipping both the temporal filter step and the FIX-ICA procedure. The percentage of overlapped time points between the DVARS and DI masks from temporal scrubbing is 9.81% of the total time points. The DVARS-based scrubbing (red circle) produced significant effects on the local ($t = 10.79$; $p = 0$; $d = 0.71$; df = 468) and long-range ($t = 2.93$; $p = 0.0039$; $d = 0.19$ df = 468) rsfMRI connectivity compared to control group. The local ($t = 8.21$; $p = 0$; $d = 0.54$; df = 468) and long-range ($t = 3.18$; $p = 0.0026$; $d = 0.21$ df = 468) scrubbing effect of the DVARS-based scrubbing on rsfMRI connectivity diminished with retaining the high DI volumes. Note that the DVARS-based and DI-based scrubbing generated almost the same local connectivity changes, which, we hypothesize, might be associated with the remaining artifacts in the rsfMRI signals since we skipped the temporal filter step and FIX-ICA procedure. The colored circles represent the averaged connectivity changes every 6 mm. The local (ROI pairs with a distance between 13 and 49 mm, gray box) and long-range (ROI pairs with a distance between 125 and 161 mm, gray box) rsfMRI connectivity changes are summarized as a bar plot and shown as an inset in the top region. The error bar represents SEM across subjects. The asterisks represent the level of significance: **, $0.001 < p \leq 0.01$, and ***, $p \leq 0.001$.

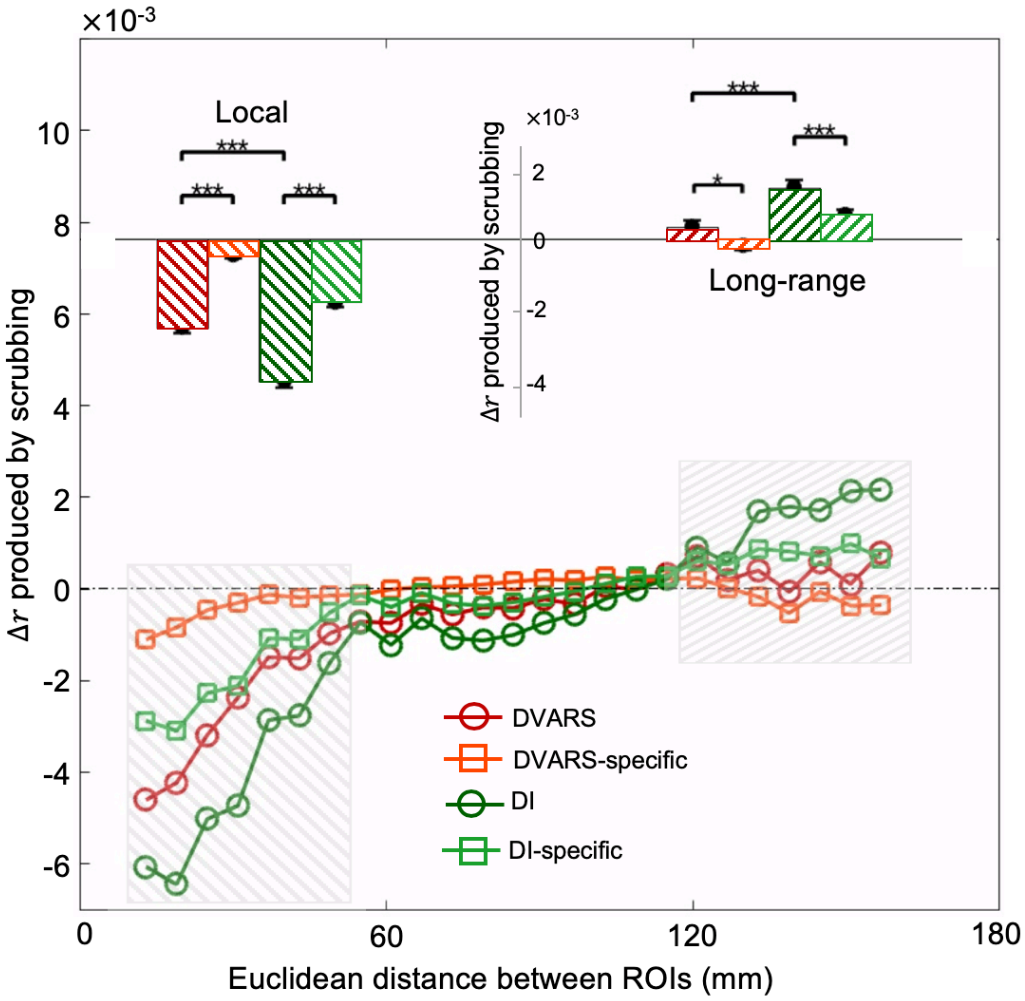


Fig. S13: Compared with the DVARS not convolved with an HRF in Fig. 4, here we investigated the scrubbing effect of DVARS and DI using DVARS convolved with the HRF. The percentage of overlapped time points between the DVARS and DI masks from temporal scrubbing is 9.54% of the total time points. The DVARS-based scrubbing (red circle) produced significant effects on the local ($t = 14.05$; $p = 0$; $d = 0.91$; df = 468) and non-significant effects on the long-range ($t = 1.30$; $p = 0.22$; $d = 0.084$; df = 468) rsfMRI connectivity compared to control group. The local scrubbing effect of the DVARS-based scrubbing on rsfMRI connectivity diminished with retaining the high DI volumes ($t = 10.41$; $p = 0$; $d = 0.67$; df = 468). The scrubbing of convolving the DVARS with the HRF produced larger local effect than that of the DVARS in Fig. 4. We hypothesize that the DVARS convolved with the HRF might have more overlapped time points with DI since the DVARS detects both motion and arousal-related changes and there is a significant delay between the motion and DI peaks (Fig. 6). The colored circles represent the averaged connectivity changes every 6 mm. The local (ROI pairs with a distance between 13 and 49 mm, gray box) and long-range (ROI pairs with a distance between 125 and 161 mm, gray box) rsfMRI connectivity changes are summarized as a bar plot and shown as an inset in the top region. The error bar represents SEM across subjects. The asterisks represent the level of significance: *, $0.01 < p \leq 0.05$, and ***, $p \leq 0.001$.

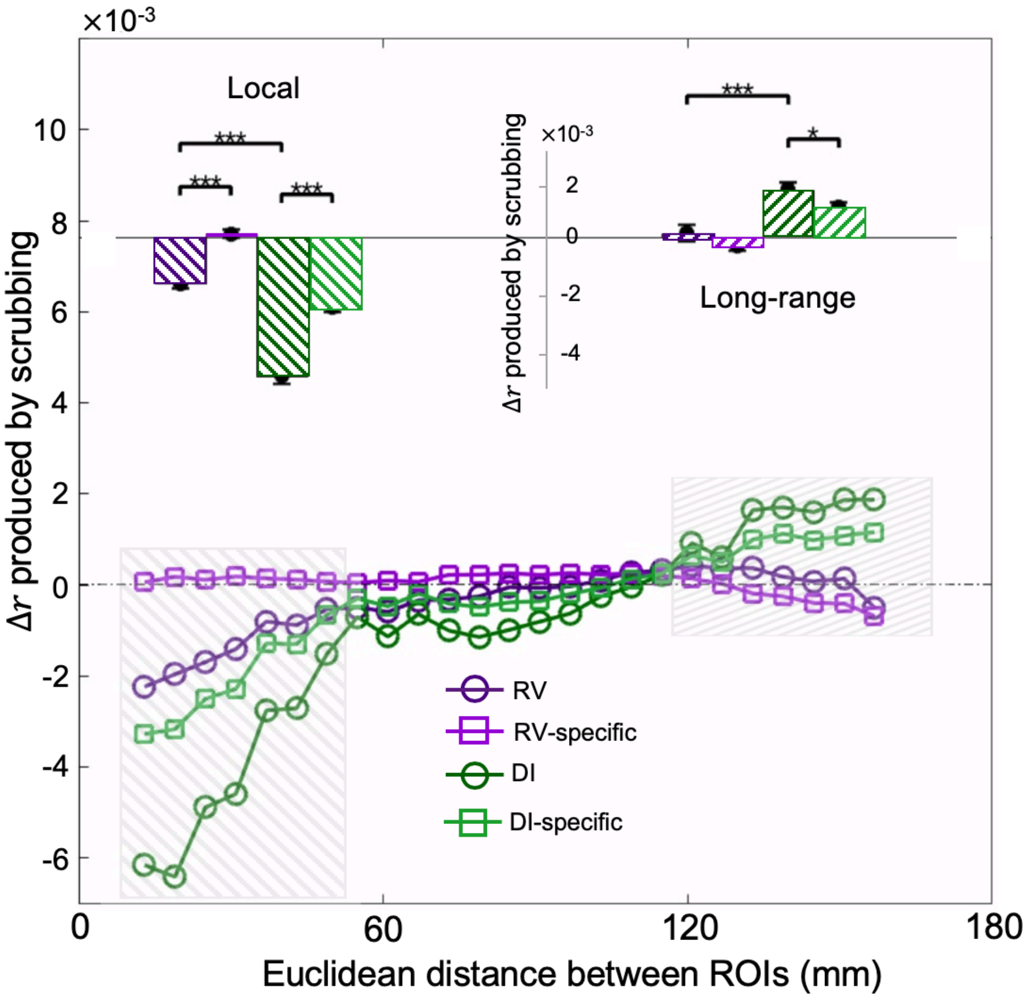


Fig. S14: Compared with the respiratory volume (RV) not convolved with a respiratory response function (RRF) in Fig. 5, here we investigated the scrubbing effect of RV and DI using RV convolved with the RRF. The percentage of overlapped time points between the RV and DI masks from temporal scrubbing is 9.73% of the total time points. The RV-based scrubbing produced significant effects on local ($t = 8.17$; $p = 2.78\text{x}10^{-15}$; $d = 0.54$; df = 406) and non-significant effects on long-range ($t = 0.14$; $p = 0.89$; $d = 0.010$; df = 406) rsfMRI connectivity compared to the control group. The local scrubbing effect of the RV-based scrubbing on rsfMRI connectivity diminished with retaining the high DI volumes ($t = 7.29$; $p = 1.09 \text{ x } 10^{-5}$; $d = 0.21$; df = 406). The scrubbing of convolving the RV with the RRF produced larger effect than that of the RV (Fig. 5). We hypothesize that the RV convolved with the RRF might have more overlapped time points with DI given that there is a significant delay between the RV and DI peaks (Fig. S20). The colored circles/squares represent the averaged connectivity changes every 6 mm. The local (ROI pairs with a distance between 13 and 49 mm, gray box) and long-range (ROI pairs with a distance between 125 and 161 mm, gray box) rsfMRI connectivity changes are summarized as a bar plot and shown as an inset in the top region. The error bar represents SEM across subjects. The asterisks represent the level of significance: *, $0.01 < p \leq 0.05$, and ***, $p \leq 0.001$.

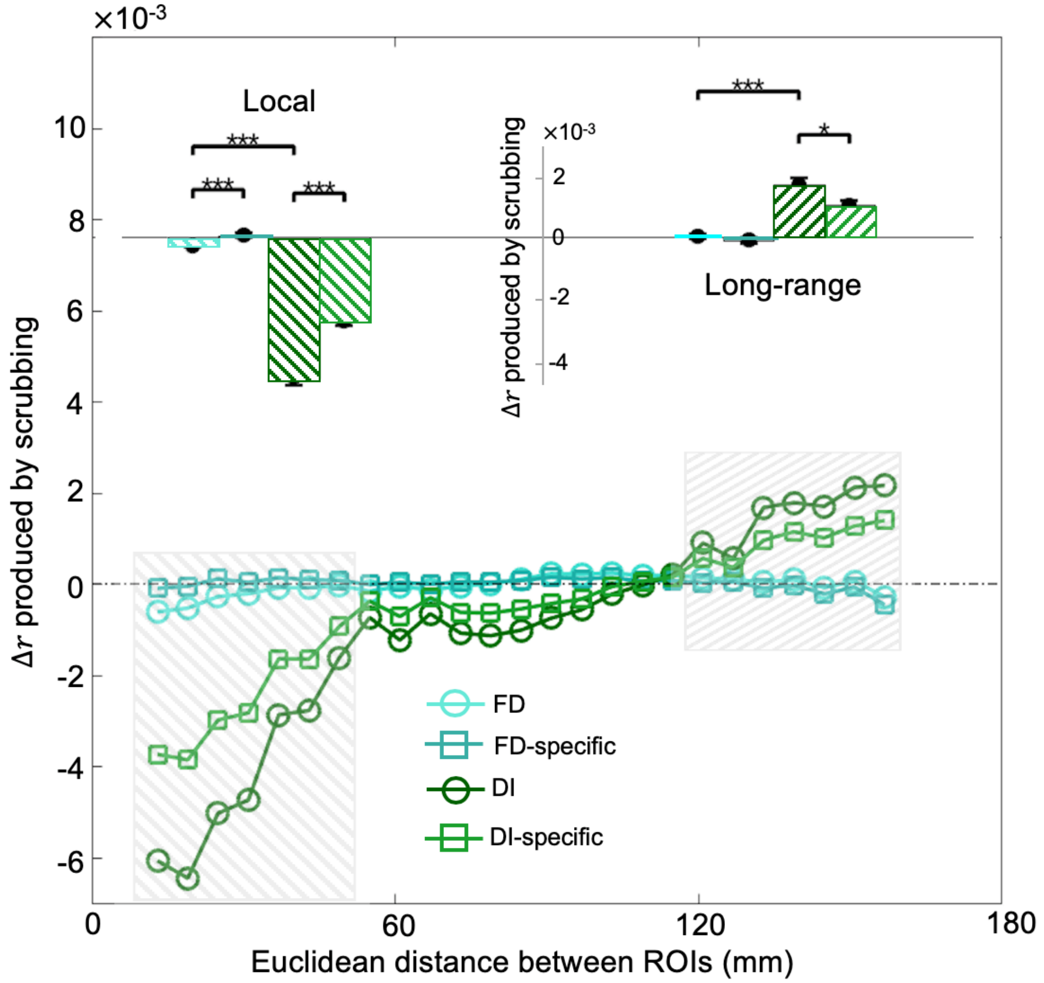


Fig. S15: The effect of the FD-based scrubbing on rsfMRI connectivity diminished with retaining the high DI volumes. The percentage of overlapped time points between the FD and DI masks is 7.16% of the total time points. The FD-based scrubbing (cyan circle) produced significant effects on the local ($t$ = 3.59; $p$ = 4.07x10$^{-4}$; $d$ = 0.23; df = 468) and non-significant effects on the long-range ($t$ = 0.99; $p$ = 0.32; $d$ = 0.064; df = 468) rsfMRI connectivity compared to the control group. The local scrubbing effect of the FD-based scrubbing on rsfMRI connectivity diminished with retaining the high DI volumes ($t$ = 3.72; $p$ = 0; $d$ = 0.24; df = 468). Thus, the effect of FD-based scrubbing on rsfMRI connectivity is attributed to time points with high DI values, which are associated with arousal modulations. The colored circles/squares represent the averaged connectivity changes every 6 mm. The local (ROI pairs with a distance between 13 and 49 mm, gray box) and long-range (ROI pairs with a distance between 125 and 161 mm, gray box) rsfMRI connectivity change are summarized as a bar plot and shown as an inset in the top region. The error bar represents SEM across subjects. The asterisks represent the level of significance: *, $0.01 < p \leq 0.05$ and ***, $p \leq 0.001$.

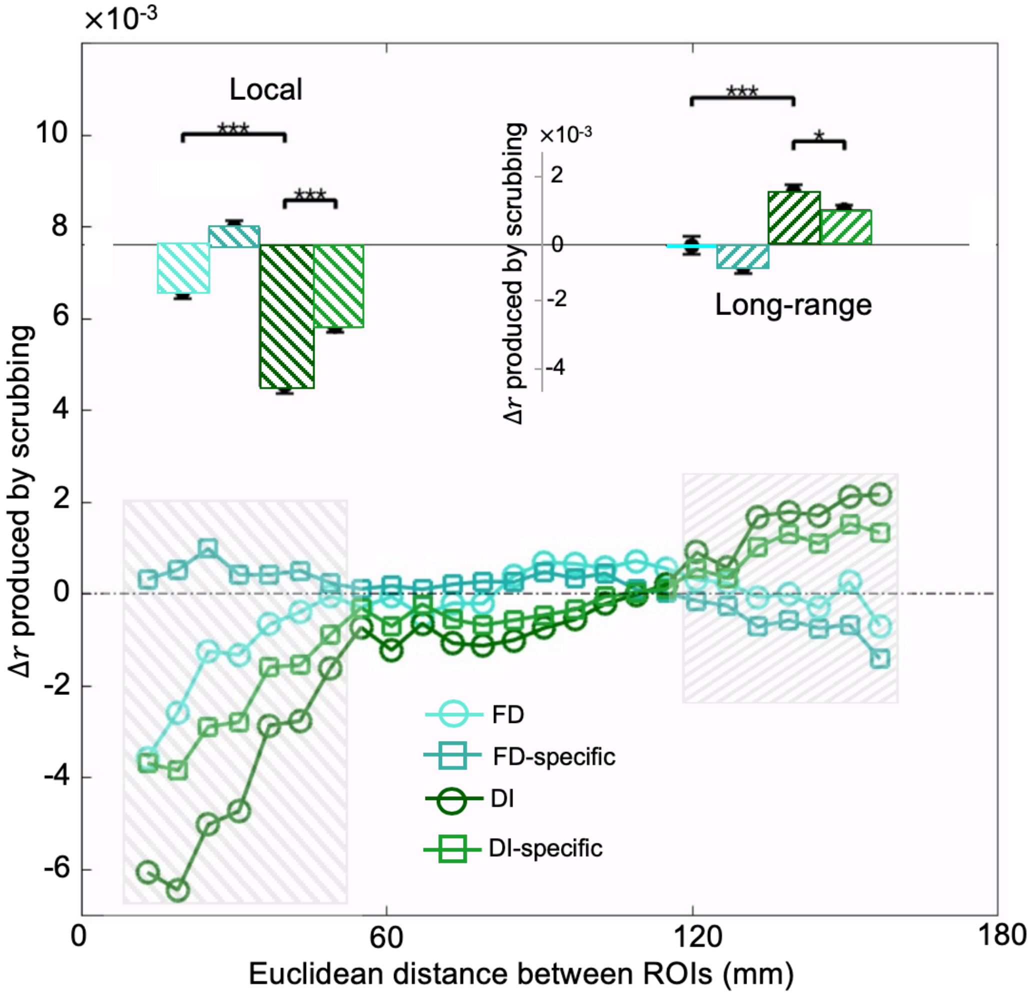


Fig. S16: Compared with the FD not convolved with a hemodynamic response function (HRF) in Fig. S15, here we investigated the scrubbing effect of FD and DI using FD convolved with the HRF. The percentage of overlapped time points between the FD and DI masks from temporal scrubbing is 8.14% of the total time points. The FD-based scrubbing (cyan circle) produced significant effects on the local ($t = 6.01$; $p = 0$; $d = 0.39$; df = 468) and non-significant effects on the long-range ($t = 0.54$; $p = 0.59$; $d = 0.036$; df = 468) rsfMRI connectivity compared to the control group. The local scrubbing effect of the FD-based scrubbing on rsfMRI connectivity diminished with retaining the high DI volumes ($t = 6.49$; $p = 0$; $d = 0.42$; df = 468). The scrubbing of convolving the FD with the HRF produced larger effect on the local connectivity than that of the FD in Fig. S15. We hypothesize that the FD convolved with the HRF might have more overlapped time points with DI given that there is a significant delay between the FD and DI peaks (Fig. 6). The colored circles represent the averaged connectivity changes every 6 mm. The local (ROI pairs with a distance between 13 and 49 mm, gray box) and long-range (ROI pairs with a distance between 125 and 161 mm, gray box) rsfMRI connectivity changes are summarized as a bar plot and shown as an inset in the top region. The error bar represents SEM across subjects. The asterisks represent the level of significance: *, $0.01 < p \leq 0.05$, and ***, $p \leq 0.001$.

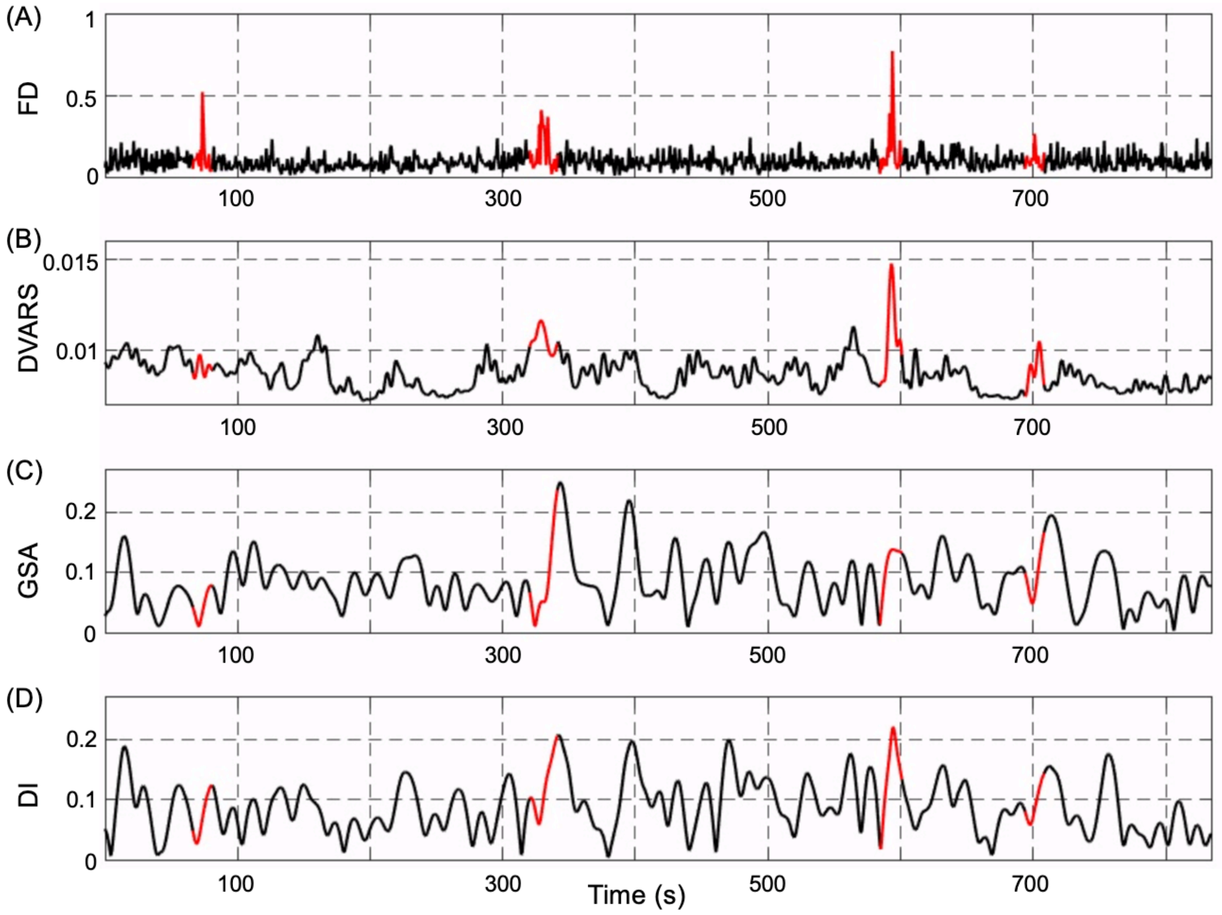


Fig. S17: Time courses of FD (A), DVARS (B), GSA (C), and DI (D) from a typical subject. FD spikes co-occur with DVARS peaks but are followed by GSA and DI peaks with significant delay. Twenty-one time points centering on FD spikes are marked as red color for all of the time courses.

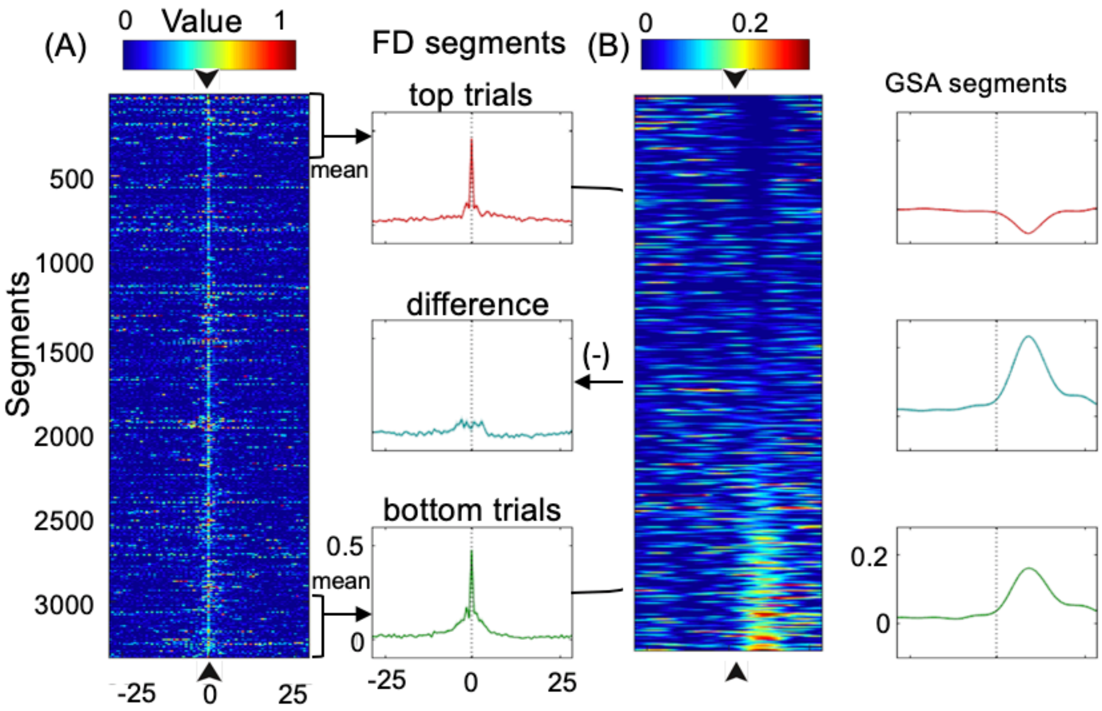


Fig. S18: Temporal relationships among FD and GSA. Segments of FD (A) and GSA (B) were extracted with respect to the 3331 identified FD spikes (time 0) and sorted according to the values of GSA at 9.36 seconds. The top (red) and bottom (green) 400 segments were averaged and their differences (cyan) were also calculated. The shadow represents regions within one SEM. The black arrows and dash lines indicate time zero.

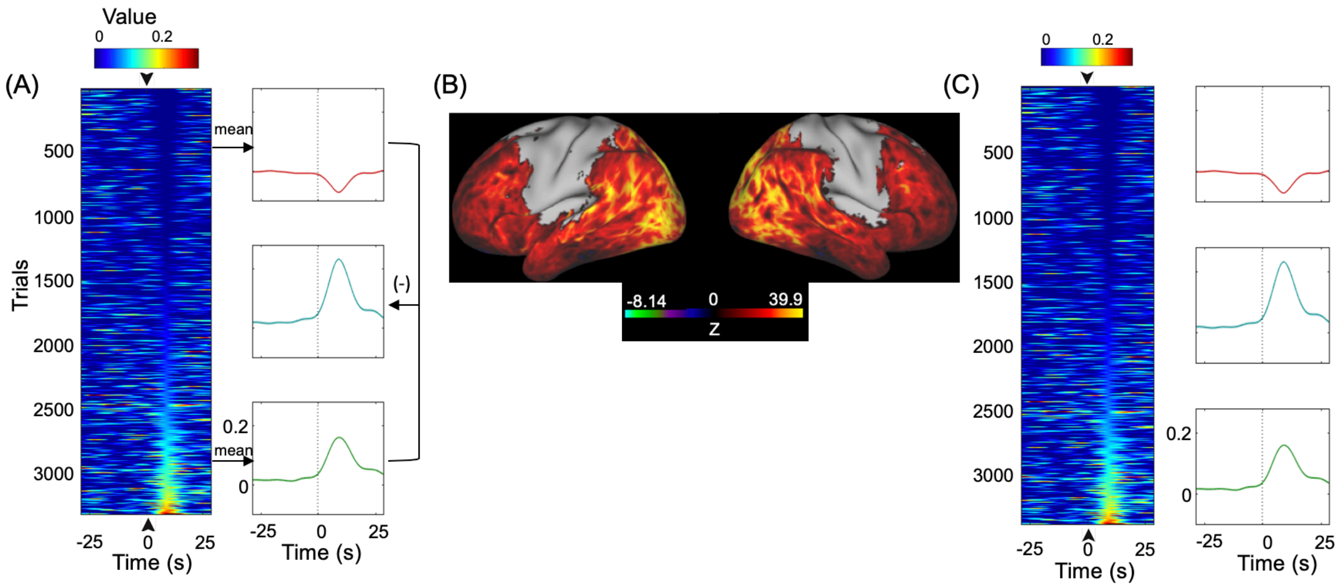


Fig. S19: DI peaks following FD spikes are not caused by sensorimotor activations related to head motion. We calculated a second version of DI, i.e., DI_noSM, from the global co-activation pattern excluding sensorimotor regions (B). Segments of DI (A) and DI_noSM (C) were extracted respectively with respect to the FD spikes and sorted by the DI values at 9.36 seconds. The top (red) and bottom (green) 400 segments were averaged and their difference (cyan) was also shown. The shadow represents regions within one SEM. The black and dash lines arrow indicates time zero. (B) The global co-activation pattern excluding the sensorimotor regions.

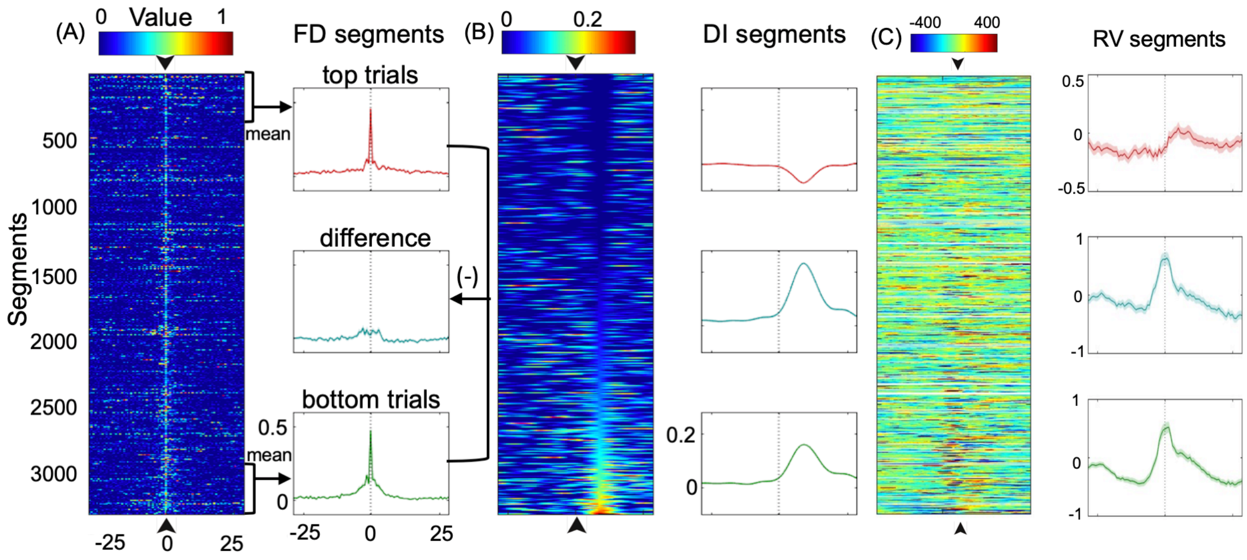


Fig. S20: Temporal relationships among FD, DI and RV. Segments of FD (A), DI (B) and RV (C) were extracted with respect to the 3331 identified FD spikes and sorted according to the DI values at 9.36 seconds. The top (red) and bottom (green) 400 segments were averaged and their differences (cyan) were also calculated. The shadow represents regions within one SEM. The black arrows and dash lines indicate time zero. Note that physiological data were available for 407 out of 469 subjects in HCP dataset. Thus, 441 out of 3331 (~13%) segments were missed and we plotted the missed respiration segments as white lines (right panel, C).

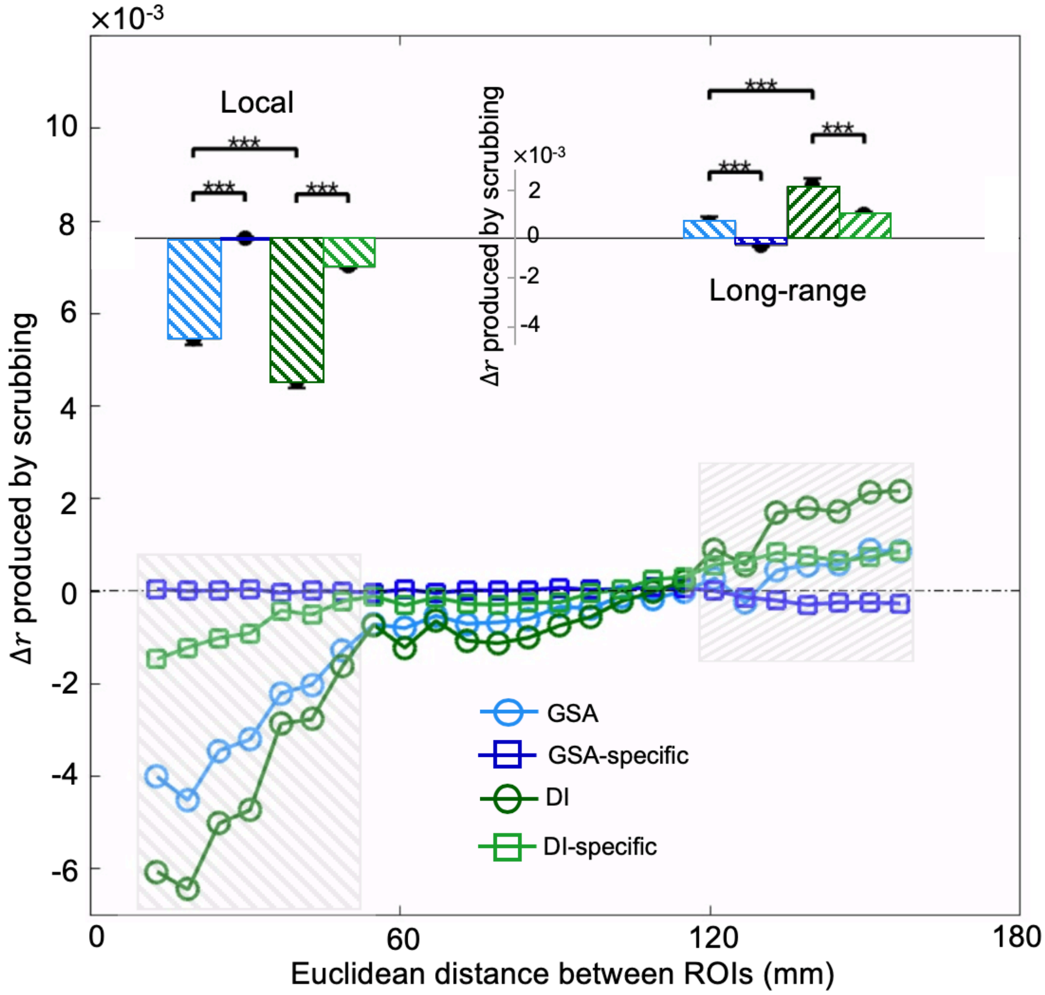


Fig. S21: The effect of the GSA-based scrubbing on rsfMRI connectivity diminished with retaining the high DI volumes. The percentage of overlapped time points between the GSA and DI masks is 19.08% of 25% the total time points. The GSA-based scrubbing (blue circle) produced significant effects on the local ($t$ = 15.37; $p$ = 0; $d$ = 1.02; df = 468) and non-significant effects on the long-range ($t$ = 1.85; $p$ = 0.073; $d$ = 0.12; df = 468) rsfMRI connectivity compared to control group. In contrast, the DI-based scrubbing (green circle) produced significant effects on both the local ($t$ = 18.39; $p$ = 0; $d$ = 1.18; df = 468) and long-range ($t$ = 6.80; $p$ = 0; $d$ = 0.46; df = 468) rsfMRI connectivity compared to control group. Moreover, the DI-based scrubbing generated larger effect on both the local ($t$ = 4.32; $p = 3.50\text{x}10^{-5}$; $d$ = 0.22; df = 468) and long-range ($t$ = 4.01; $p = 9.22\text{x}10^{-5}$; $d$ = 0.28; df = 468) connectivity changes than the GSA-based scrubbing. The local ($t$ = 15.56; $p$ = 0; $d$ = 1.04; df = 468) and long-range ($t$ = 3.39; $p = 4.1\text{x}10^{-4}$; $d$ = 0.22; df = 468) scrubbing effect of the GSA-based scrubbing on rsfMRI connectivity diminished with retaining the high DI volumes. To summarize, the DI-based scrubbing outperformed the GSA-based scrubbing given that the global co-activation template used to calculate DI incorporated the spatial information of arousal-related fMRI changes. The colored circles/squares represent the averaged connectivity changes every 6 mm. The local (ROI pairs with a distance between 13 and 49 mm, gray box) and long-range (ROI pairs with a distance between 125 and 161 mm, gray box) rsfMRI connectivity changes are summarized as a bar plot and shown as an inset in the top region. The error bar represents SEM across subjects. The asterisks represent the level of significance: ***, $p \leq 0.001$.

Table S1. Pearson's correlation of all alert-related behavioral measures with FD, DVARS, GSA and DI (degree of freedom: 467; FDR-corrected $p$ values less than 0.05 are highlighted)

| Behavioral measures | FD | | DVARS | | GSA | | DI | |
|---|---|---|---|---|---|---|---|---|
| | Pearson $r$ | FDR-corrected $p$ values | Pearson $r$ | FDR-corrected $p$ values | Pearson $r$ | FDR-corrected $p$ values | Pearson $r$ | FDR-corrected $p$ values |
| PSQI_Score | 0.14 | **0.027** | 0.060 | 0.30 | 0.045 | 0.47 | 0.063 | 0.30 |
| Bed time | 0.13 | 0.053 | 0.16 | **0.0057** | 0.21 | **0.00014** | 0.21 | **0.00013** |
| Amount of sleep | -0.12 | 0.074 | -0.030 | 0.65 | -0.089 | 0.15 | -0.14 | **0.027** |
| Minutes to fall asleep | 0.080 | 0.22 | 0.0092 | 0.89 | 0.10 | 0.11 | 0.12 | 0.067 |
| Get up time | 0.020 | 0.70 | 0.093 | 0.14 | 0.12 | 0.07 | 0.093 | 0.14 |
| Cannot get to sleep within 30 minutes | 0.12 | 0.070 | 0.055 | 0.36 | 0.062 | 0.31 | 0.073 | 0.25 |
| Wake up during sleep | 0.11 | 0.081 | -0.0045 | 0.93 | -0.0083 | 0.89 | 0.023 | 0.73 |
| Get up to use bathroom | 0.070 | 0.28 | -0.031 | 0.64 | -0.031 | 0.64 | -0.047 | 0.45 |
| Breathe uncomfortably | 0.16 | **0.0070** | 0.097 | 0.12 | 0.062 | 0.31 | 0.090 | 0.15 |
| Snore loudly | 0.23 | **0.000030** | 0.17 | **0.0031** | 0.014 | 0.84 | 0.049 | 0.43 |
| Feel too cold | 0.10 | 0.11 | 0.070 | 0.26 | 0.015 | 0.83 | 0.046 | 0.46 |
| Feel too hot | 0.14 | **0.027** | 0.044 | 0.48 | -0.10 | 0.11 | -0.056 | 0.35 |
| Have bad dream | 0.078 | 0.23 | 0.0087 | 0.89 | 0.068 | 0.27 | 0.072 | 0.25 |
| Have pain | 0.077 | 0.23 | 0.076 | 0.24 | -0.061 | 0.31 | -0.043 | 0.49 |
| Other sleep troubles | -0.0050 | 0.93 | -0.072 | 0.25 | -0.097 | 0.12 | -0.11 | 0.10 |
| Overall sleep quality | 0.075 | 0.25 | 0.0042 | 0.93 | 0.057 | 0.35 | 0.10 | 0.11 |
| How often taken sleep medicine | -0.041 | 0.51 | 0.016 | 0.83 | -0.039 | 0.53 | -0.067 | 0.28 |
| Trouble staying awake | 0.10 | 0.11 | 0.061 | 0.31 | 0.078 | 0.23 | 0.095 | 0.13 |
| Trouble keeping up enthusiasm | 0.074 | 0.25 | 0.072 | 0.25 | -0.029 | 0.65 | 0.0098 | 0.89 |
| Have bed partner or roommate | -0.042 | 0.50 | -0.11 | 0.089 | -0.095 | 0.13 | -0.070 | 0.26 |
